# Energy-guided combinatorial co-optimization of antibody affinity and stability

**DOI:** 10.1101/2025.11.26.690765

**Authors:** Ariel Tennenhouse, Adva Mechaly, Tony Pham, Robert J. Oldham, Jake Henry, Dorry X. Zhao, Thomas M. Moon, Isabel G. Elliott, Omri Porat, Jinny Kim, Yael Fridmann Sirkis, Sveta Gaiduk, Itay Talpir, Jamie Bird, Christopher Domalewski, Pei-Hsuan Chu, Andrew B. Dippel, Rebecca Croasdale-Wood, Daniel Christ, Gilad Kaplan, Mark S. Cragg, Sarel J. Fleishman

**Affiliations:** Department of Biomolecular Sciences, Weizmann Institute of Science, Rehovot, Israel; Department of Infectious Diseases, Israel Institute for Biological Research, 7410001 Ness-Ziona, Israel; Biologics Engineering, R&D, AstraZeneca, Gaithersburg, MD, USA; Antibody & Vaccine Group, Center for Cancer Immunology, School of Cancer Sciences, Faculty of Medicine, Southampton General Hospital, University of Southampton, Southampton, SO16 6YD, UK; Garvan Institute of Medical Research, Sydney, NSW 2010, Australia; Protein Analysis Unit, Department of Life Sciences Core Facilities (LSCF), Weizmann Institute of Science, Rehovot, Israel; Biologics Engineering, R&D, AstraZeneca, Cambridge, UK

## Abstract

Affinity maturation is an essential process in antibody engineering. Although powerful, it is iterative, time-consuming, and can result in trade-offs, where affinity is gained at the cost of important properties, such as specificity and stability. We present a scalable strategy, called LAffAb, that starts from a crystallographic structure of the antibody-antigen complex and introduces combinations of mutations to optimize its energy. A combinatorial library comprising 7,000 variants with up to nine mutations results in gains of up to 30-fold in affinity while co-optimizing stability. Surprisingly, the library does not converge on a single solution, instead favoring diverse variants with a high mutational load. Small-scale screening of 10 designs against a potential drug target results in an order-of-magnitude improvement in affinity while maintaining high developability. We also apply LAffAb to improve the developability of a therapeutic antibody without degrading affinity. We envision that LAffAb can be used to design stable, specific, and high-affinity binders and to improve our understanding of sequence, structure, and function relationships in antibodies.

## Introduction

Molecular recognition is critical for many biological functions, with antibodies providing a prime example. Due to their tremendous versatility and compatibility with the human immune system, antibodies have been applied as therapeutics against pathological conditions ranging from cancer to neurological and autoimmune diseases, and are ubiquitous reagents in all branches of biomedical research^1–3^.

Binding affinity is one of several parameters that require optimization for antibodies to be effective. Their binding surfaces are mostly formed by six solvent-exposed loops called the complementarity-determining regions (CDRs). The animal immune system improves antigen-binding affinity by introducing random mutations, primarily in the CDRs, and selecting the highest-affinity variants for further randomization in an iterative process called affinity maturation^4^. *In vitro* affinity maturation is currently the dominant strategy for engineering higher-affinity antibodies, often improving affinity by orders of magnitude relative to the parental molecule^5^. But this process is time-consuming and may result in unacceptable trade-offs in which high affinity is gained only by compromising other properties, such as stability and specificity^6–8^. In the context of developing an antibody therapeutic, in addition to affinity, other biophysical properties, including colloidal stability, low self-association, and minimal nonspecific binding (collectively termed “developability”) are essential^9^. Antibody engineering is therefore a complex, multidimensional problem that requires balancing multiple biophysical traits that may be mutually exclusive. Beyond these practical concerns, the fact that a critical step in developing antibodies for basic and applied biomedical research still relies on random processes reflects significant gaps in our understanding and control over antibody sequence, structure, and function.

The primary challenge for reliable design in the CDRs is the complexity of their structure. Antibodies often bind antigenic surfaces lacking defined secondary structure and presenting both polar and hydrophobic groups^10^. To form high-affinity and specific interactions, the CDRs must form a complementary surface that typically lacks secondary structure elements. To maintain these conformations even in the absence of the antigen, the CDRs are stabilized by an intricate network of polar and nonpolar interactions within and between the CDRs and with the framework of the antibody variable fragment (Fv)^11,12^. Given this structural complexity, design calculations must carefully weigh the potential gain from forming new interactions with the antigen against the risks of removing crucial interactions for antibody stability and foldability.

Several computational strategies have been developed to address the limitations of antibody affinity maturation^13–15^. Due to the low likelihood of obtaining beneficial multipoint mutants, these approaches have relied on iterative mutational screening of single mutations. In one study, for instance, single-point mutations that improved electrostatic complementarity with the antigen were experimentally screened, and favorable ones were combined to improve affinity^13^. Large language models have also been used to improve affinity. Here too, however, single-point mutations were experimentally screened before generating combinations^14^, and contemporary language models cannot accurately predict antibody developability without fine-tuning on experimental data^16,17^. For instance, a recent application of an ensemble of language models was used to design mutations to co-optimize antibody expression and affinity, but only 19 of 132 screened single-point mutations were beneficial, requiring experimental screening of single and double mutants to reach multi-point mutants that ultimately exhibited less than threefold improvement in binding signal^18^. Furthermore, language models have generally been successful when applied to antibodies for which large sets of mutational effects were available^14,15^, and simulations suggest that successful training of AI-driven methods for affinity prediction may require two orders of magnitude more affinity measurements than are available in publicly accessible databases^19^. Thus, design methods have been restricted to iterative screening procedures^20^.

Here, we report a non-iterative structure- and energy-based approach, called LAffAb (**L**ibraries of **Aff**ined **A**nti**b**odies), to design multipoint mutants in antibody CDRs that optimize native-state energy. We first tested the reliability of LAffAb by designing and screening a small combinatorial library of CDR mutants. Deep sequencing showed that most designed mutations were allowed, and many combinatorial designs were preferred over the parental antibody. Applied at small scale to an antibody of interest for therapeutic development, LAffAb improved affinity by 30-fold without compromising developability. Finally, we targeted the therapeutic antibody Urelumab, and many designs maintained affinity while improving thermostability and reducing undesirable self-association. Unlike conventional computational and experimental methods, LAffAb does not iterate mutagenesis and screening and produces large and diverse sets of gain-of-function mutants without compromising stability.

## Results

### 1. Mutation tolerance mapping in the CDRs

We start the design process by mapping the mutational tolerance of the CDRs based on a crystallographic structure of the antibody-antigen complex and Rosetta atomistic calculations (Fig. 1, steps 1 and 2). Rosetta design calculations optimize native-state energy and have been successfully used to optimize biologics for several properties, including stability, expression, and binding^21–27^. We start by computationally mutating each CDR position to Ala and flagging positions in which mutations increase system energy (Fig. 1, step 3). Atomistic energy functions tend to penalize buried polar and electrostatic interactions due to desolvation^28^, favoring hydrophobic ones in their place; however, buried polar interactions are crucial for foldability, and mutating them may lead to misfolding and dysfunction^29–31^. To mitigate the risk of mutating crucial buried polar residues, we also flag positions as immutable if they contribute substantially to hydrogen-bonding energy (Fig. 1, step 4). Following these restrictions, the design workflow follows the rationale of the FuncLib approach^32^, which has been successfully used to optimize obligatory and non-obligatory binding surfaces^22,26,30^ and enzyme active sites^32,33^. Briefly, we exclude mutations that are rare in a multiple-sequence alignment (MSA) of antibodies (calculated separately for the heavy and light chains) in all regions except CDR H3. Because of its diversity, MSAs of H3 are unlikely to be reliable; instead, we use previously inferred per-position residue frequencies^34^. High-frequency mutations are individually modeled in the context of the antigen-bound structure and scored in Rosetta, and mutations that substantially increase system energy are triaged (Fig. 1, step 5). Our hypothesis is that the pre-filtering steps based on atomistic calculations and observed frequencies remove mutations that destabilize the antibody or deform its backbone, limiting the effects of long-range epistasis between mutations^35^. As an optional step, a few mutations are included in the design if visual inspection determines that they maintain or improve molecular interactions (Supp. Tables 1-3).

**Figure 1:**
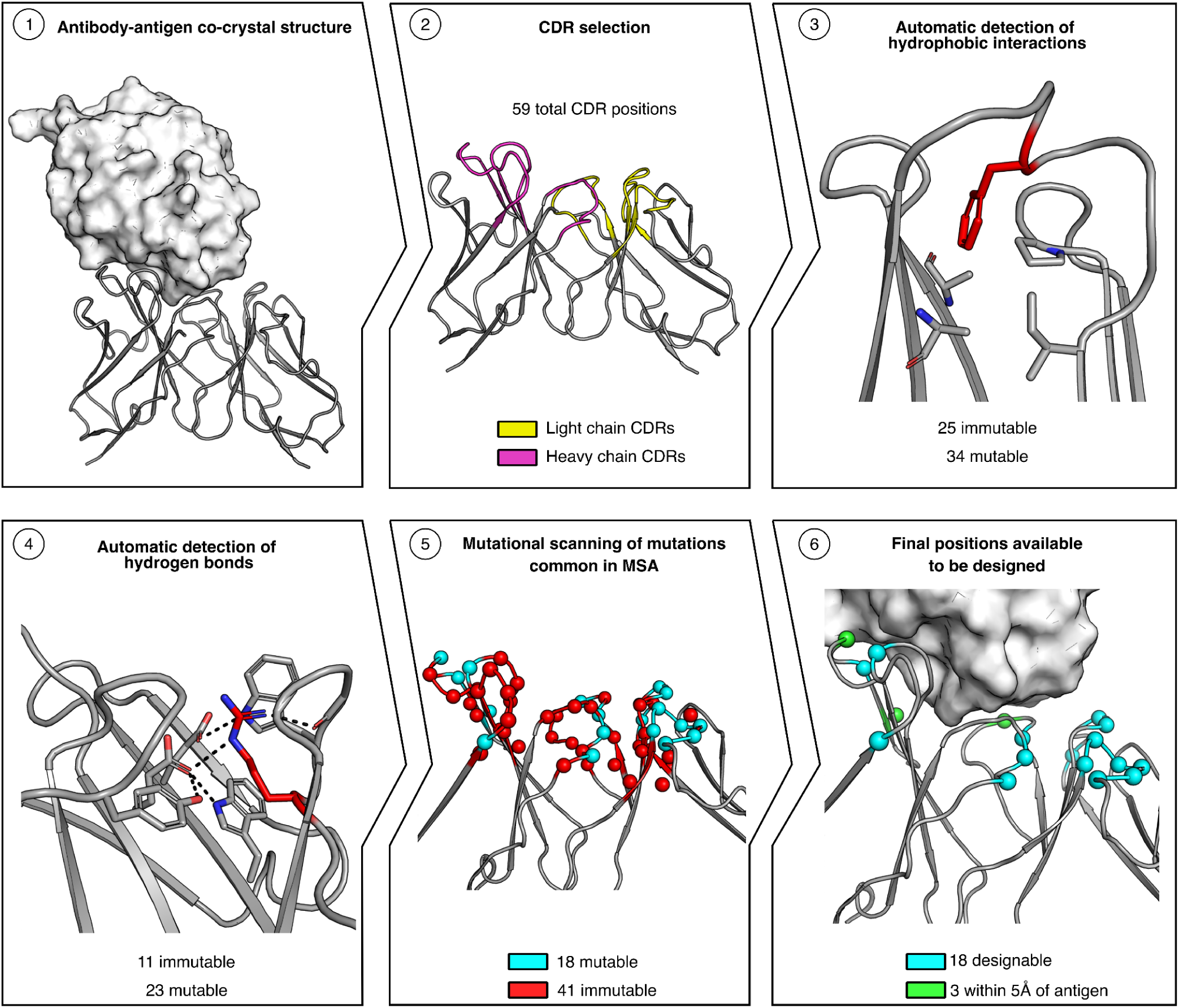
Computational mutation tolerance mapping in the antibody Fv. LAffAb begins from a co-crystal structure of an antibody-antigen complex (Step 1). CDR boundaries are automatically determined (Step 2; Supp. Table 4). Atomistic design calculations flag all positions in which mutation to Ala substantially worsens total or hydrogen-bonding energy (Steps 3 and 4, respectively). Then, mutations allowed by phylogenetic screening are modeled at each allowed CDR position, and those that substantially compromise stability are eliminated (Step 5). Mutations were mostly allowed in the periphery of the interface; only three positions in direct contact with the antigen were mutable (Step 6). Models were generated based on the co-crystal structure of D44.1 and HEWL (PDB ID 1MLC ref. 36). In steps 5 and 6, Cα atoms of indicated residues are shown as spheres. HEWL shown as a white surface map.

To test the reliability of LAffAb, we designed a medium-sized library of variants based on the murine anti-hen egg white lysozyme (HEWL) D44.1 antibody^36^, starting from a crystallographic structure of the antigen-binding fragment (Fab) bound to its antigen (PDB ID 1MLC). D44.1 has served as a model system for studying affinity maturation for decades^13,22,37^, allowing us to benchmark LAffAb against other approaches. Moreover, this is a high-affinity interaction (*K_D_* of approx. 5 nM)^21,38^, and D44.1 underwent animal affinity maturation, making it representative of a typical antibody-engineering task.

Mutation tolerance mapping showed that only 18 of 59 CDR positions could be mutated without destabilizing the CDRs or the antigen-antibody complex. The majority of those (15) did not form direct interactions with the antigen (Fig. 1, step 6). D44.1, therefore, presents a case in which most CDR positions are unlikely to tolerate mutations, particularly in the most critical antigen-binding surface. Nevertheless, we hypothesized that mutations in solvent-exposed parts of the CDRs may indirectly contribute to affinity by stabilizing the conformations of the CDRs, thereby promoting the antigen-bound state as observed in the crystallographic structure.

### 2. Substantial affinity improvements by one-shot library design

We designed a focused combinatorial library starting from the tolerated sequence space of D44.1. The sequence space of all tolerated mutations at the 18 positions was too large to enumerate (over 1 billion mutation combinations). For computational tractability, we first removed mutations in positions far from the binding surface and ones that introduced noncomplementary charges with the antigen (Supp. Table 1). Only two of the 17 designed point mutations (Asn92_L_Ala and Ser56_H_Val) were shared with a previous computational design study dedicated to improving the affinity of D44.1^13^. A significant challenge for combinatorial-library design is that mutations that are individually tolerated may be destabilizing in combination, in a phenomenon known as negative epistasis^35^. To limit this risk while also minimizing design calculations, we modeled all combinatorial mutants within each CDR separately from the other CDRs (except for L2 and H3, which are proximal and were enumerated in combination), amounting to only 115 atomistic design calculations. Each calculation modeled the combinatorial mutant and iterated sidechain packing and constrained sidechain and backbone minimization. Finally, the low-energy designs from all CDRs were combined, yielding a combinatorial library of 6,912 variants. We verified that >99% of these designs exhibited more favorable Rosetta energy than the parental antibody (Fig. 2A), suggesting that the library would be enriched for stabilized, potentially higher-affinity variants.

**Figure 2:**
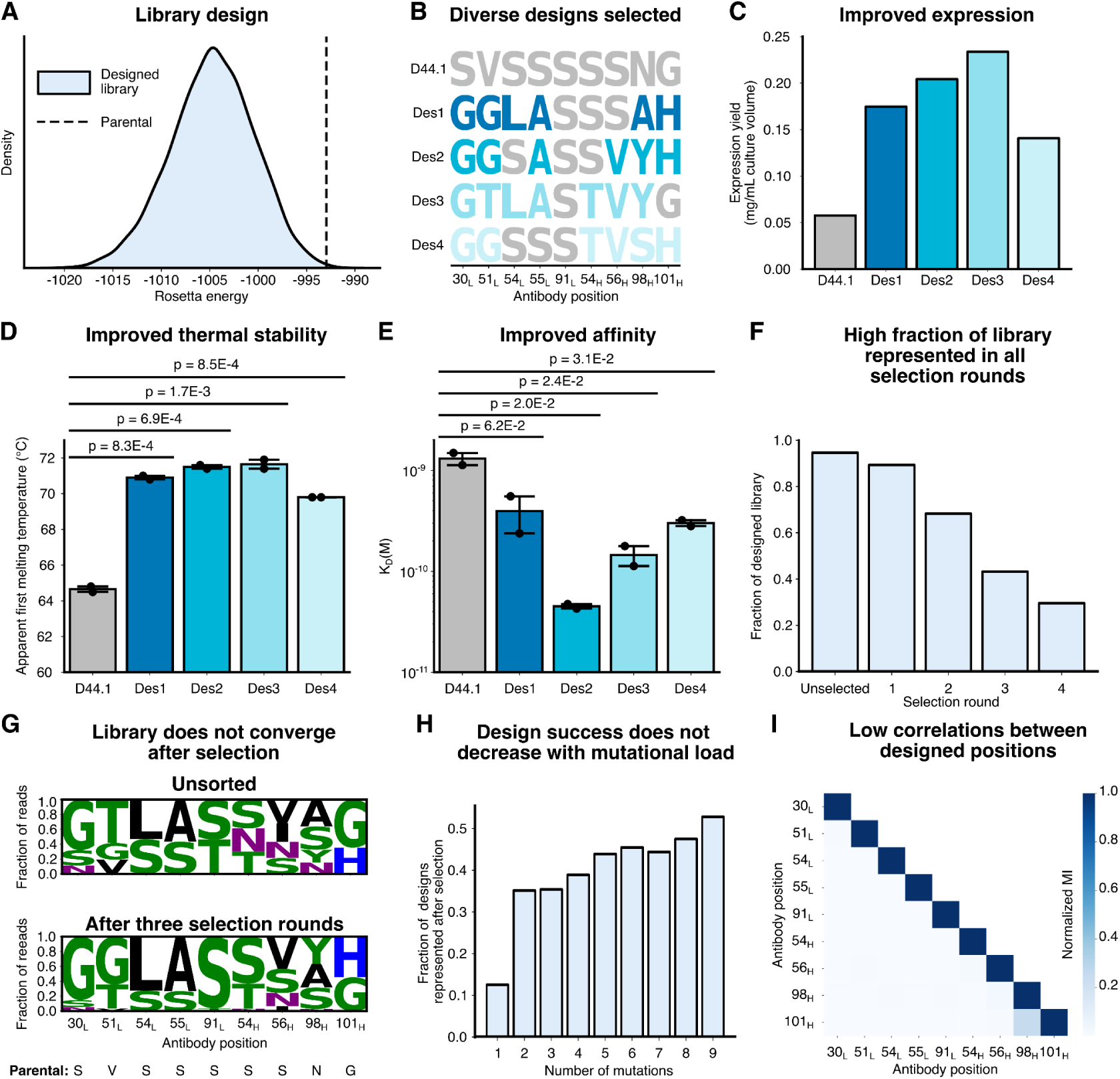
Design and selection of D44.1 variants that co-optimize affinity and stability. (**A**) Rosetta energy distribution of the designed library comprising 6,912 variants. Over 99% of the designed library exhibits improved Rosetta energy compared to D44.1. (**B**) Mutations at each designed position in D44.1. (**C**) Expression yield (corrected by purity) of antibodies formatted as mouse IgG2a. Based on a single measurement. (**D**) Apparent first melting temperatures measured by nano differential scanning fluorimetry (nanoDSF). (**E**) Dissociation constants (*K_D_*) measured by SPR. (**D, E**) Bars and error bars represent the mean and standard deviation of two technical replicates (measurements plotted as dots). *P*-values calculated using a Student’s *t-*test. (**F**) Fraction of the designed library present in at least one read after each phage-selection round. (**G**) Sequence logoplots of the population as a function of the selection round. Parental identities noted on bottom. (**H**) Fraction of the designed library observed in selection round 3 by number of mutations. (**I**) Mutual information analysis of positions in selected population.

We assembled the library into single-chain variable fragments (scFvs) from DNA oligos using GGAssembler^39^ and screened them for binding using phage display^40^. After four rounds of panning, 22 individual colonies were sequenced. Surprisingly, we isolated 19 unique designs, none of which was the parental D44.1 (Supp. Table 5). This suggested that despite its small size, the library comprised multiple solutions that were superior to the parent. Furthermore, the selected designs exhibited 4-8 mutations from D44.1 (Supp. Fig. 1A), an unusually large number of mutations in single-shot affinity-maturation campaigns. The designs were assayed for binding to HEWL using monoclonal phage ELISA, and all showed a higher binding signal than D44.1, with some exhibiting dramatic improvements (Supp. Fig. 1B).

We selected four distinct designs with 6-7 mutations from D44.1 for expression as full-length murine IgG2a antibodies (Fig. 2B). After single-step protein A purification, three exhibited purity matching or exceeding that of D44.1 (Supp. Fig. 2A, B). All four exhibited improved expression yields in Chinese hamster ovary (CHO) cells of as much as fourfold (Fig. 2C) and increases in thermal stability relative to D44.1 (Fig. 2D, Supp. Fig. 2C). Finally, surface plasmon resonance (SPR) experiments revealed large gains in affinity, with design Des2 exhibiting over 30-fold improvement, reaching 50 pM affinity (Fig. 2E, Supp. Fig. 2D). We concluded that one-shot LAffAb calculations produced diverse variants with many CDR mutations relative to conventional single-shot antibody design and engineering processes, and that the designs exhibited large improvements in affinity together with gains in stability.

### 3. Distinct high-affinity solutions revealed by deep sequencing

We applied deep sequencing analysis to understand why so many different multipoint mutants were selected from the designed library (Supp. Table 6). Even after four selection rounds, 30% of the designed sequences exhibited at least one read (Fig. 2F), though the parental D44.1 was purged by the third round. After the third selection round, most designed mutations were allowed, and some were vastly preferred relative to the parental identity (Fig. 2G, Supp. Table 7). Deep sequencing further showed that the fraction of designs with at least one read after three selection rounds showed no decrease as a function of mutational load (Fig. 2H). We used the deep sequencing data to test our hypothesis that the designed mutations are minimally epistatic. Indeed, a mutual-information analysis revealed very low correlations in the selected population between all positions except two structurally adjacent ones in the heavy chain (98 and 101), which were weakly correlated (Fig. 2I). The results strikingly demonstrate that selecting tolerated point mutations according to energy and structural criteria dramatically reduces the risk of negative epistasis in combinations of individually favorable mutations. They further suggest that the space and diversity of beneficial multipoint mutants in CDRs may be very large.

Given that a large fraction of the library apparently exhibited improved affinity, we sought to understand which individual mutations failed to improve affinity. We calculated the frequency of each mutation in the selected versus unselected populations compared to the parental identity (Supp. Table 7). Ten of the 16 designed single mutations had a normalized enrichment greater than one, suggesting they were beneficial compared to the parental identity. Three others had a normalized enrichment over 0.5, suggesting they were not strongly purged. Only three mutations were strongly counter-selected: Ser91_L_Thr, Ser54_H_Asn, and Ser56_H_Ile (Fig. 2G, Supp. Table 7), a very low fraction of deleterious mutations relative to contemporary methods for antibody design^18^.

We evaluated the Rosetta system energy of all single-point mutations, but the energies did not correlate with normalized enrichment (Supp. Fig. 3A), excluding this computed metric as a predictor. We therefore analyzed finer Rosetta energy-based filters for clues to explain the negative impact of the counter-selected mutations. Ser91_L_Thr was in a position flagged by the computational alanine scan, because the mutation weakens a hydrogen bond to light chain His34 (Fig. 3A, right panel). We nevertheless added this deleterious mutation to test how sensitive the selection process is to conservative mutations. Structural analysis suggested that the other two deleterious mutations each introduced strain to the local structure that is relieved in modeling through a subtle backbone change (0.4 and 0.6 Å between Cα atoms of the parental structure and design model of the single-point mutant) or a change in the conformation of the antigen (Fig. 3A, left panels). By contrast, the 13 favorable or neutral mutations did not abrogate hydrogen bonds, and all mutations that were enriched relative to the parental identity (Supp. Table 7) did not alter the backbone by more than 0.4 Å (Supp. Table 8, Supp. Fig. 3B). Surprisingly, whereas Ser56_H_Ile was strongly counterselected, Ser56_H_Val was tolerated. The dramatic influence of a single methyl emphasizes the need for accurate atomistic design methods (compare Fig. 3A bottom left panel and Fig. 3B left panel). Additionally, we visually inspected the design model of the highest-affinity design. Two neighboring small-to-large mutations, Asn98_H_Tyr, and Gly101_H_His, may stabilize the backbone of CDR H3 by forming additional interactions with nearby positions and by reducing the desolvation penalty of the partly buried Asn in the parental D44.1 (Fig. 3B bottom right panel). Interestingly, Gly101_H_His was not strongly favored as an individual mutation according to the deep sequencing results (Fig. 2G, Supp. Table 7), highlighting the potential of combining several weakly enhancing mutations to generate large gains in affinity.

**Figure 3:**
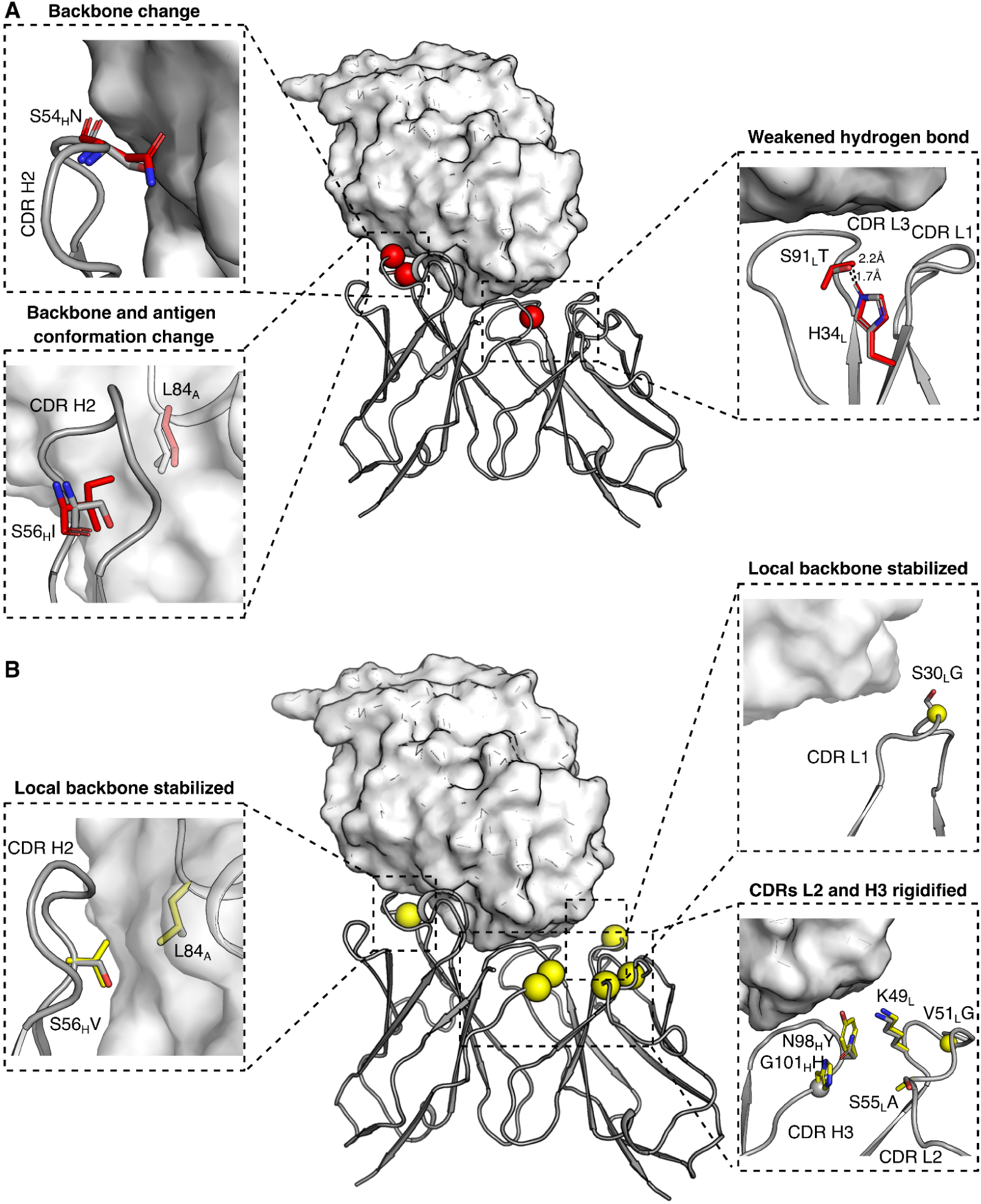
Structural underpinnings of favorable and deleterious mutations. (**A**) Ser91_L_Thr (right panel) weakens a hydrogen bond with His34_L_. Ser54_H_Asn (top left panel) shifts the position of the main chain backbone atoms, potentially introducing strain into CDR H2. Similarly, Ser56_H_Ile (bottom-left panel) changes the position of the main chain backbone atoms. Furthermore, the mutation forces a change in the conformation of Leu84 in HEWL to accommodate the larger residue. (**B**) Ser56_H_Val likely stabilizes CDR H2 because of its β-branched side chain (left panel). A cluster of small-to-large mutations likely stabilizes CDR H3, particularly by improving interactions with Lys49 of CDR L2. (**A and B**) The middle structure shows the Rosetta-relaxed model structure of the D44.1 Fv in complex with HEWL shown as a surface map. The Cɑ atoms of the mutations are shown in red (deleterious mutations) or yellow (favorable) spheres in the middle panel. Inset panels show the Rosetta-relaxed model in gray and mutations in colors. Spheres in the inset panels represent glycines. Subscript L denotes light chain, subscript H denotes heavy chain, and subscript A denotes antigen (HEWL).

We concluded that, in addition to selecting mutations that exhibit low energy, CDR design calculations should avoid mutations predicted to distort the local backbone structure, cause a change in the conformation of the antigen, or remove internally stabilizing hydrogen bonds. These structure- and energy-based rules can be applied to any antibody-antigen co-crystal complex to dramatically enrich beneficial combinations of mutations. Indeed, with the benefit of hindsight, had we excluded the three deleterious mutations, the size of the library would have been reduced to 1,728 variants, of which 1,381 (80%) were observed in deep sequencing after three selection rounds. We note that including the visually selected Ser91_L_Thr mutation in the library allowed us to verify the importance of the LAffAb pre-filtering steps. As this was the only mutation we added by visual inspection, the success of LAffAb is mostly attributed to the automated design process.

### 4. Large gains in affinity through low-throughput screening

We next tested whether small-scale application of LAffAb to an unrelated antibody-antigen complex would also improve affinity. We applied the design process described above to the 6G08 antibody, which targets the sole inhibitory crystallizable fragment (Fc) receptor, Fc gamma Receptor IIb (FcγRIIb)^41^, an attractive drug target due to its role in regulating the immune response^41^. However, for some applications, antibodies targeting FcγRIIb must also avoid binding to FcγRIIa, a key activating FcγR with 93% sequence identity to FcγRIIb in the extracellular domain^42^ (Fig. 4A). Antibodies fulfilling these criteria were discovered using phage display, and in co-administration experiments, these increased the effectiveness of target-cell-deleting antibodies in mouse models^41^. Likely due to the harsh requirement to avoid binding to the homologous FcγRIIa during selection, the resulting affinities were low, and one specific antibody, 6G08, had particularly low affinity. 6G08, therefore, presented a stringent test of the ability of LAffAb to improve affinity without compromising specificity.

**Figure 4:**
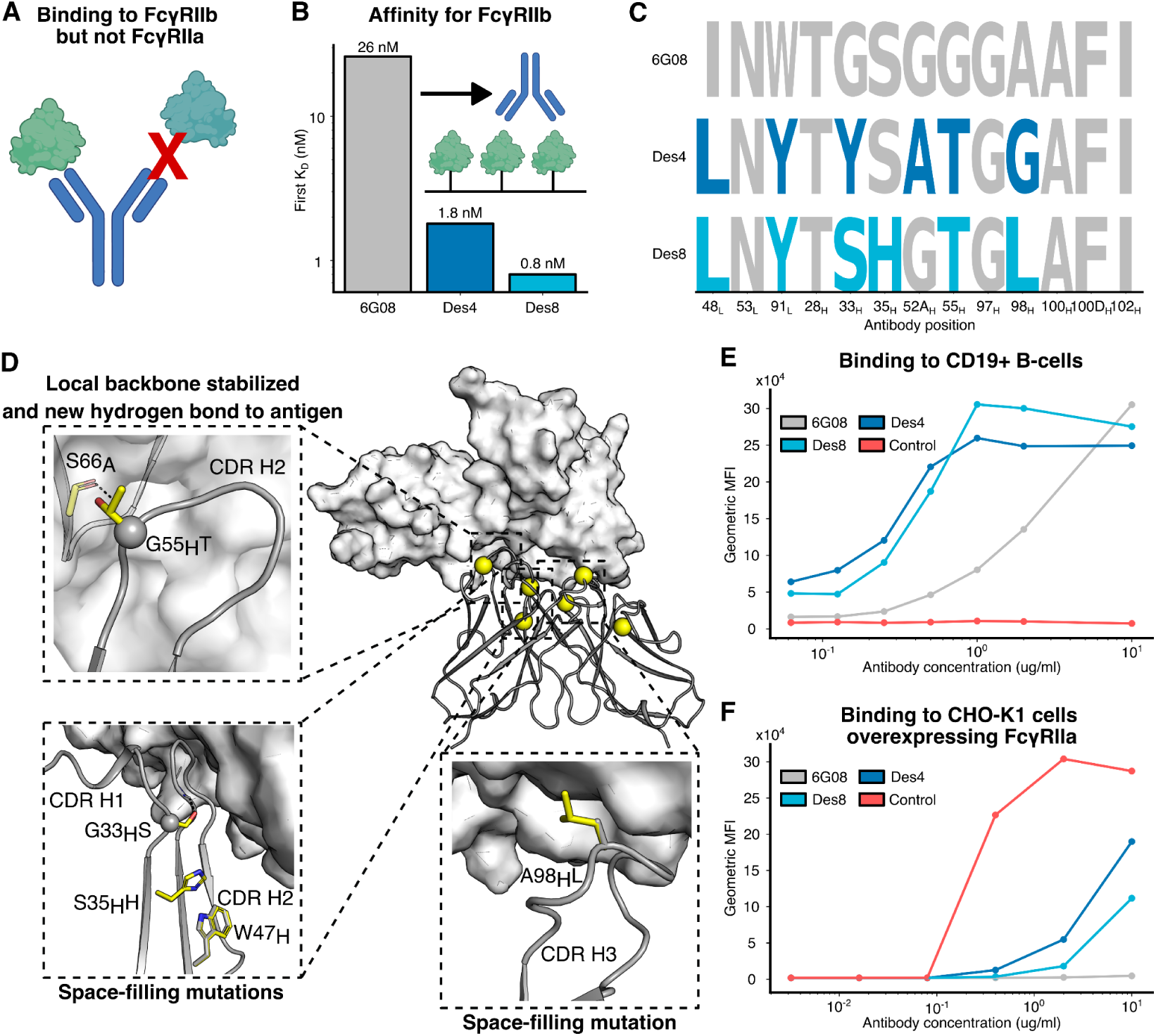
Improving affinity through small-scale experimental screening of antibody 6G08. (**A**) The desired outcome of the design process is high-affinity binding to FcγRIIb and not its close homolog FcγRIIa. Scheme made with BioRender. (**B**) Dissociation constants measured using SPR with FcγRIIb immobilized and antibodies flown over the chip. Inset scheme made using BioRender. Based on a single measurement. (**C**) Mutations at each designed position in 6G08 and the two successful designs. (**D**) Structural reasoning for affinity-enhancing mutations in Des8. Gly55_H_Thr likely stabilizes CDR H2 because of its β-branched side chain and makes a hydrogen bond with the backbone of Ser66 in FcγRIIb (top left panel). Two small-to-large mutations likely fill a hole in the antibody-antigen interface. Ser35_H_His likely strengthens a hydrogen bond to Trp47_H_ (bottom left panel). Ala98_H_Leu fills a cavity in the antibody-antigen interface (right panel). The middle structure shows the Rosetta-relaxed model structure of the 6G08 Fv in complex with FcγRIIb shown as a surface map. The Cɑ atoms of the mutations in Des8 are shown in yellow spheres in the middle panel. Inset panels show the Rosetta-relaxed model in gray and mutations in yellow. Spheres in the inset panels represent glycines. Subscript L denotes light chain, subscript H denotes heavy chain, and subscript A denotes antigen (FcγRIIb). (**E**) Binding to B-cells isolated from PBMCs. B-cells naturally express FcγRIIb. (**F**) Binding to FcγRIIa using CHO-K1 cells overexpressing FcγRIIa. (**E and F**) Two independent experiments were performed using either directly conjugated reagents or detection with a fluorescently labeled secondary antibody. The same trend was observed in both experiments. The results using secondary detection are presented.

We applied computational mutation tolerance mapping to the co-crystal structure of 6G08 bound to FcγRIIb^43^ (PDB ID 5OCC). Unlike the D44.1-HEWL complex, our calculations suggested many stabilizing mutations in the antibody-antigen interface. We speculate that the reason for the difference between the two antibodies is that 6G08 had not undergone affinity maturation^41^. We enumerated all combinations of point mutations in the CDRs and selected ten designs with six mutations relative to 6G08 and 3-10 mutations relative to one another (Supp. Table 9). Some positions were flagged by the computational alanine scan but were included because they improved the system and/or hydrogen-bonding energy (Supp. Table 2). We predicted the immunogenicity of designs using the Next-Generation IEDB Tools webserver^44^. The ten designs added at most one class II peptide compared to 6G08, and the number of predicted immunogenic peptides was comparable to Pembrolizumab (Supp. Table 10).

Eight of the ten designs bound FcγRIIb, and two exhibited much improved affinity of up to 30-fold (Fig. 4B, Supp. Fig. 4). The two designs had four mutations relative to each other, suggesting that LAffAb was again able to produce distinct high-affinity solutions (Fig. 4C). Structural modeling showed that mutations in the two successful designs were predicted to form new interactions with the antigen, fill cavities in the antibody-antigen interface, and stabilize the CDR backbones (Fig. 4D). One mutation (Trp91_L_Tyr) that was present in both successful designs was added by visual inspection; however, the mutation was predicted by Rosetta to be beneficial and was not flagged for hydrogen bonding. Ser35_H_His was present in one successful design and added during visual inspection, but it was also in many unsuccessful designs. The other two mutations added during visual inspection were each present in only one, unsuccessful design. Thus, only one or two of the six mutations in the successful designs were introduced manually, and both were favored by energy calculations.

To verify that the designs function in a physiological setting, they were assayed for binding to FcγRIIb-expressing CD19+ B cells within isolated peripheral blood mononuclear cells (PBMCs) by flow cytometry (Fig. 4E), and both showed greatly improved binding relative to 6G08. Additionally, the designs were tested for binding to CHO-K1 cells overexpressing the homologous off-target FcγRIIa (Fig. 4F), and both showed only weak binding, verifying that specificity was mostly maintained. Furthermore, both designs saturated binding to the CD19+ B cells at 1 μg/mL, and at this concentration, both showed very weak binding to the CHO-K1 cells, suggesting there is a significant window in which the designs can be given without detectable binding to FcγRIIa, even when FcγRIIa is overexpressed.

In addition to affinity, other developability characteristics are crucial for a therapeutic lead. We performed a panel of developability assays on 6G08 and the two successful designs, revealing that the designs maintained the favorable expression yield, thermal stability, low self-association, and low polyspecificity of the parental antibody (Supp. Fig. 5, Supp. Table 12). Thus, the affinity of 6G08 for its intended target was enhanced without compromising developability, while maintaining specificity toward a very close, evolutionarily conserved homolog of the target.

### 5. Developability improvement of a therapeutic antibody

The large space of high-affinity variants LAffAb generates presents an opportunity to select designs predicted to maintain affinity while improving other desirable properties. We applied LAffAb to Urelumab, a high-affinity therapeutic antibody that targets the immune receptor 4-1BB (K_D_ 8 nM) and is known to exhibit high reversible self-association^9^. Self-association and nonspecific binding are correlated with low and high isoelectric point (pI) in the Fv, respectively, and the pI of Urelumab is 5.9, outside the desirable range (6.5-8.5)^45^. We selected designs with pI in the desirable range (Supp. Table 13), and as a control, tested designs that passed the initial filtering but exhibited high Rosetta system energy. In this case, no mutations were added during visual inspection. All designs had at most two additional peptides predicted to bind MHC Class II and had fewer predicted peptides than Pembrolizumab (Supp. Table 14).

We tested 38 low- and 14 high-energy designs for binding to 4-1BB and using several developability assays measuring aggregation, nonspecific binding, and reversible self-association (Supp. Table 15). The low-energy designs had significantly higher thermal stability than the high-energy ones, but most other criteria were indistinguishable among these two groups (Supp. Fig. 6). 24 low-energy designs maintained affinity within twofold of Urelumab (Fig. 5A, Supp. Fig. 7), and nearly all (36) showed improvements in thermal stability (Fig. 5B). We classified each design as passing or failing developability based on resistance to aggregation, reversible self-association, and nonspecific binding (Supp. Fig. 8). 17 passed all assays, and six of these exhibited comparable affinity to Urelumab (Fig. 5A). The six high-affinity designs that passed all developability tests were diverse, comprising 4-6 mutations from Urelumab and at least two mutations relative to one another (Fig. 5C). Thus, LAffAb can generate multiple diverse solutions that improve developability while maintaining affinity.

**Figure 5:**
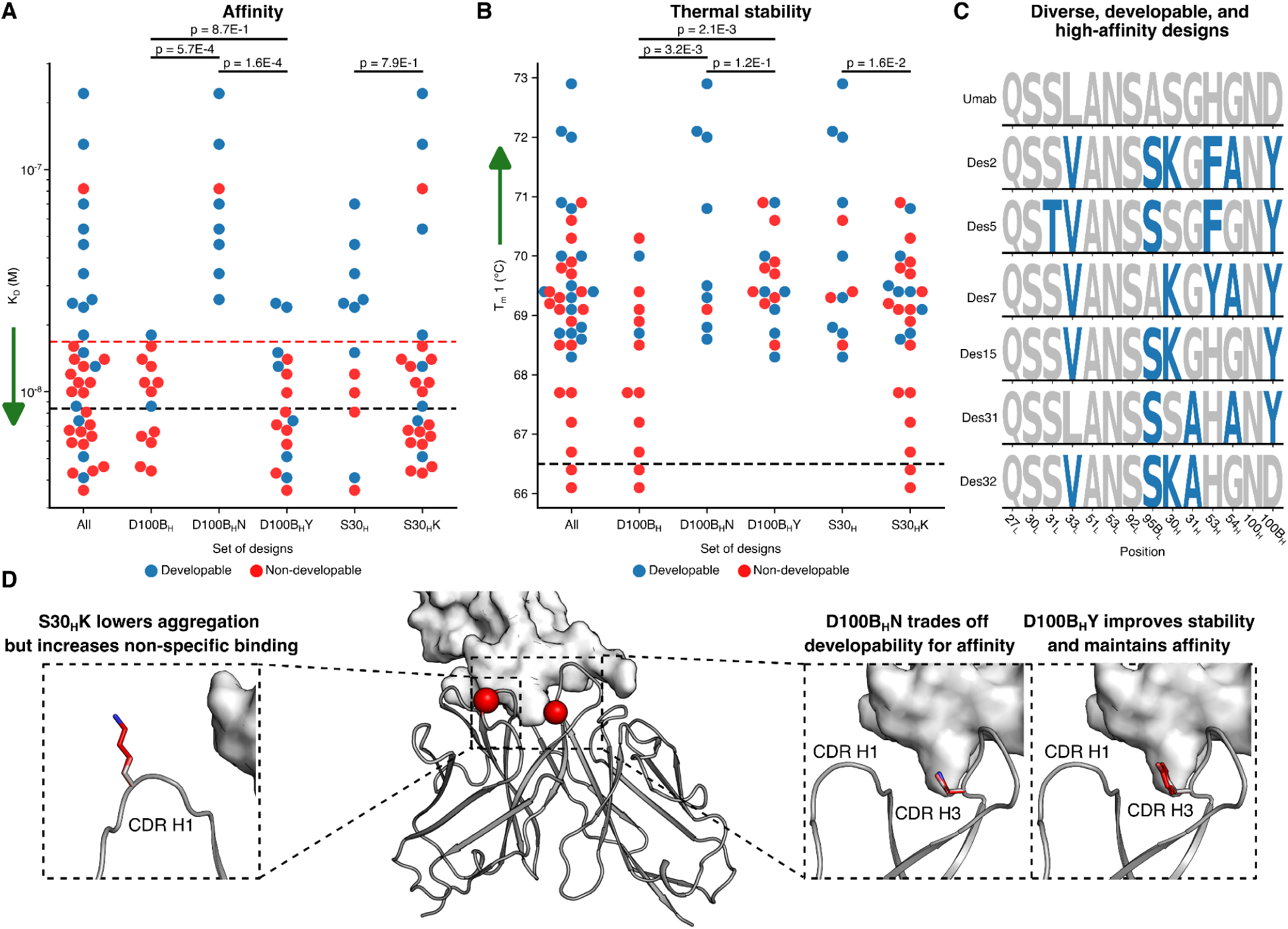
Improving developability through medium-scale experimental screening of Urelumab. (**A**) Affinities of different design subsets measured by SPR. Red dashed line represents a twofold decrease in the affinity of Urelumab. Sensorgrams shown in Supp. Fig. 7. (**B**) Thermal stability of different design subsets measured by nano differential scanning fluorimetry (nanoDSF). (**A,B**) Black dashed line represents the value of Urelumab. *P*-values calculated using a Student’s *t-*test. Dots represent the means of three technical repeats. Green arrows represent the preferred direction in each assay. (**C**) Mutations at each designed position in Urelumab (Umab) and the six high-affinity, developable designs. (**D**) Structural reasoning for developability effects of selected mutations. The middle structure shows the Rosetta-relaxed model structure of the Urelumab Fv in complex with 4-1BB shown as a surface map. The Cɑ atoms of the selected mutations are shown in red spheres in the middle panel. Inset panels show the Rosetta-relaxed model in gray and mutations in red. (**A, B, and D**) Subscript H denotes heavy chain.

We analyzed the individual mutations that comprised the designs to identify mutations leading to improvements or losses in each assay. Asp100_B_Asn in the heavy chain significantly improved stability and developability compared to designs containing the parental Asp but compromised affinity (Fig. 5 A, B, and D). Asp100_B_Tyr improved stability and developability without compromising affinity (Fig. 5 A, B, and D). In both cases, the improvements in developability were driven by significant reduction in reversible self-association and nonspecific binding (Supp. Figs. 8B and C).

Ser30Lys in the heavy chain significantly improved aggregation resistance while maintaining affinity (Fig. 5A and Supp. Fig 8A) but increased reversible self-association (Supp. Fig. 8B) and nonspecific binding (Supp. Figs. 8C, D). Despite these liabilities in designs containing Lys30, four of six high-affinity and developable designs contained Lys30 (Fig. 5C). This suggests that other mutations compensated for the nonspecific binding and self-association seen with Lys30 while maintaining the benefits in aggregation resistance.

## Discussion

Mutation tolerance mapping is key to the success of LAffAb, ensuring that the mutations are individually allowed and retain the CDR backbone conformation. Maintaining the backbone conformation is crucial to enable combining multiple mutations that do not antagonize one another, but is complicated by the irregularity and sensitivity of the CDRs to mutations. The careful mutation tolerance mapping in LAffAb identifies a “smooth” local sequence space where mutations are mostly structurally and energetically compatible and complementary with one another, limiting negative epistasis^35^. Thus, LAffAb is distinguished from previous natural, experimental, and computational maturation methods that prioritize single mutations without explicitly assessing their impact on the structure. Furthermore, by optimizing the energy of the antigen-bound structure subject to constraints that favor foldability, our approach, in effect, co-optimizes affinity and stability.

Our results demonstrate that CDR positions may contribute to affinity, even if they do not interact directly with the antigen. This is important because interfacial positions are typically less tolerant to mutation. The three case studies presented very different scenarios: in D44.1, almost no antigen-binding position was mutable; in 6G08, both high-affinity designs contained interfacial mutations; and in Urelumab, many designs contained interfacial mutations. Furthermore, we applied LAffAb to design 12 variants of the Cetuximab antibody^46^, which binds epidermal growth factor receptor (EGFR), and all designs exhibited significant decreases in binding affinity. We speculate that this failure resulted from abrogating an elaborate network of ordered water molecules surrounding CDR H3 (Supp. Fig. 9). The crystal structure we originally used for design (PDB entry: 6ARU) did not show these water molecules, whereas a higher resolution structure (PDB entry: 1YY9^46^) did. Because the atomistic calculations did not consider water molecules, designed mutations likely disrupted interactions critical for foldability. Our study thus highlights the need for accurate bound structures to predict mutational tolerance and the need for design protocols that address water-mediated interactions. It also suggests that some antibodies may be more difficult targets for affinity enhancement (i.e., Urelumab), likely due to intensive prior engineering. Even in such cases, however, optimizing the native-state energy may benefit developability. The study further highlights the remarkable structural diversity in antibody-antigen complexes and how different structures impose dramatically different mutational constraints.

For decades, antibody affinity optimization has been a critical step in antibody engineering that relied on iterative maturation processes^47^. LAffAb is the first approach to our knowledge that enables directly optimizing affinity and other important biophysical properties without iterations. Notably, our study highlights the effectiveness of using structure and energy-based criteria to generate designs and to explain their success and failure. We envision that mutational tolerance mapping can be further improved by leveraging experimental data accumulated in academia and pharma companies^48^, thereby developing general ML/AI methods based on energy criteria for improving antibody design methodology. Improved CDR design methods will be broadly applicable in antibody engineering, including in the design of improved synthetic antibody repertoires^49^. The high pace of advances in antibody modeling and design methods in recent years is encouraging that reliable and automated methods for one-shot optimization of antibody affinity and developability may be within reach.

## Methods

### Computational methods

All Rosetta design simulations used git v.d9d4d5dd3fd516db1ad41b302d147ca0ccd78abd of the Rosetta biomolecular modeling software, which is freely available to academics at http://www.rosettacommons.org. All RosettaScripts^50^ XMLs can be found at https://github.com/Fleishman-Lab/LAffAb_public.

#### Selecting a set of allowed CDR mutations

We defined the CDRs once for all antibodies in this study based on previous definitions^21,51,52^ and visual inspection. CDR definitions can be found in Supp. Table 4. For each antibody, a position-specific scoring matrix (PSSM) was computed as described previously^24^ for the light and heavy chain variable regions separately. Mutations were allowed at each CDR position outside of CDR H3 if they had a PSSM score >= −2. Within CDR H3, allowed mutations were initially identified based on per-position residue frequency data^34^. For each H3 category (“short”, “medium”, and “long” defined by Prassler et al^53^.), identities were allowed at each position if they had a frequency >= 4%. Positions labeled in Prassler et al. as “E” and “F” (“medium” CDRH3 length) or “N” and “O” (“long” CDRH3 length) were considered to be the last two residues before positions 101 and 102. All mutations were allowed in positions 101 and 102.

As a first step, the antibody Fv structure in complex with the antigen was relaxed through cycles of sidechain and harmonically restrained backbone minimization and combinatorial sidechain packing in the entire Fv (see xmls/Relax.xml). Each CDR position was then mutated to alanine and scored using the REF15^28^ score function (see xmls/alascan.xml) and the REF15 score function with the weights of all terms not related to hydrogen bonding set to zero (see xmls/alascan_Hbond.xml). Positions were considered designable if the mutation to alanine had ΔΔ*G* <=3.5 REU above the parental using the REF15 score function and <= 0.6 REU using the hydrogen bond only score function. Each mutation allowed by the PSSM or CDR H3 residue frequency data was then individually modeled and scored in Rosetta (see xmls/filterscan.xml). In conventional FuncLib calculations, the PSSM score is weighted and added to the Rosetta system energy to bias the design towards the consensus among homologous sequences^32^. Because LAffAb targets the CDRs, which are subject to strong Darwinian selection, this bias is inappropriate, and the design is not biased towards the highest-frequency identities.

Mutations were also modeled at positions flagged as making critical interactions, and individual mutations were allowed in these positions only if they maintained or improved these interactions (by visual inspection). For D44.1, only Ser91Thr in the light chain was allowed in a flagged position (Supp. Table 1). For 6G08, Trp91Tyr in the light chain, Ser35His in the heavy chain, Phe100_D_Leu in the heavy chain, and Ile102Val in the heavy chain were allowed in flagged positions (Supp. Table 2). For Urelumab, no mutations were allowed in flagged positions (Supp. Table 3).

#### Selecting combinatorial CDR mutants

Before combinatorial enumeration, completely solvent-exposed mutations, mutations far from the interface, and mutations that introduced unmatched charges were removed. (Supp. Tables 1-3). For D44.1, all combinations of allowed mutations within each CDR were enumerated. CDR L2 and CDR H3 were enumerated together because of their structural proximity. Each mutant was modeled in Rosetta with no added contribution from the PSSM score, including combinatorial sidechain packing, and the backbone and sidechains of all residues were minimized, subject to harmonic restraints on the Cɑ coordinates of the entire protein (xmls/combinatorial_enumeration.xml). Each design was labeled as positive if the all-atom energy was more favorable than that of the parental antibody and negative otherwise. Mutations were removed if the variant for that mutation had ΔΔ*G* > 1 REU.

For D44.1, an EpiNNet^54,55^ neural network model was then trained to predict which designs exhibited lower-energy than the parental antibody-antigen complex for each CDR (or L2 and H3 together). EpiNNet was used for D44.1 because we aimed to design a combinatorial library as opposed to individual designs. In short, a multi-layer perceptron classifier with a single hidden layer the size of the number of selected positions was trained on all the data. Each classifier was trained for 20,000 iterations on a one-hot encoded representation of the sequence data to classify whether a design exhibited lower energy than the parental antibody. Next, the single-point mutations were ranked according to their probability of being lower-energy than the parent. Mutations were selected for the library by iteratively adding the top-ranked mutations until the fraction of designs exhibiting lower energy than the parental antibody-antigen complex significantly dropped. All combinations of the selected mutations for each CDR were then enumerated. Each mutant was modeled in Rosetta following the same procedure.

For 6G08, all combinations of allowed mutations within the entire Fv were enumerated. Each mutant was modeled in Rosetta, including combinatorial sidechain packing, and the backbone and sidechains of all residues were minimized, subject to harmonic restraints on the Cɑ coordinates of the entire protein (see xmls/combinatorial_enumeration.xml). The designs were ranked by all-atom energy, and EpiNNet was not used in the design process. Only designs with fewer than seven mutations were considered.

For 6G08, positions in the antibody-antigen interface were defined using the InterfaceByVector Residue Selector in Rosetta on the antibody-antigen crystal structure (see xmls/interface.xml). Asn53 in the light chain and Gly50 in the heavy chain were manually removed from the description of the interface despite being included in the positions selected by InterfaceByVector. Designs were clustered so that each design had at least three interface mutations from all other designs, and the top ten clustered designs ranked by total score were selected.

For Urelumab, the pI of the Fv of each design was calculated using the ProtParam^56^ module of BioPython^57^, and only designs with pI greater than 6.5 and less than 8.5 were considered. Designs that introduce N-glycosylation sites (NX[ST] where X is not Pro) were purged. 44 positive designs and 15 negative designs were selected. For each, the designs were ranked according to Rosetta energy (with low-energy designs at the top of the list for positive designs and the bottom for negative designs). Four (one) designs with six mutations, ten (three) with five mutations, and thirty (11) with four mutations were chosen for the positive and negative pools. Within each group, designs were clustered such that there were at least three, two, and two mutations between designs. After design selection, if any positions had less than one third of designs containing the parental identity, 0.1 REU was subtracted from all designs containing the parental identity in these positions and designs were selected again. This process was iterated until all parental identities were present in at least one third of the designs. EpiNNet was not used in the Urelumab design process.

#### Analysis of Nanopore sequencing

The pod5 files output from nanopore were basecalled using the Dorado v1.0.0 duplex basecaller with the super-accurate basecalling model. The resulting basecalled reads were then filtered to remove simplex reads that were already accounted for as duplex reads using SAMtools view^58^ (v1.17). The filtered reads were then demultiplexed using Dorado demux, and the adaptors were trimmed using Dorado trim.

Each read was aligned to a FASTA sequence of each of the constant segments between designed variable regions (six constant segments total) using MiniMap2^59^ v2.30 with a minimizer kmer length of 21 (-k 21) and a minimizer window size of 11 (-w 11). The alignments were filtered to exclude unmapped reads and non-primary alignments using SAMtools view^58^ (-F 3844). PySam^58^ v0.21.0 was used to iterate through the alignments for each database and filter the reads based on several properties. The query for each alignment needed to be over 850 base pairs and less than 900 base pairs to be considered. Additionally, the query for each alignment needed to have an average PhRed^60^ score >= 20. Each alignment needed to be less than five base pairs shorter than the length of each constant segment and less than five base pairs longer than the length of each constant segment. The designed variable segments of each read were extracted based on the positions of the corresponding constant segments from the alignments. Variable segments with a high probability (>= 0.9) of having no errors based on the per-base pair PhRed scores were searched against a database of the designed variable segments, and reads with a perfect match to each designed variable segment were considered.

#### Analysis of individual mutations based on Nanopore sequencing data

The frequencies of each identity at each position at each selection round were calculated as follows:

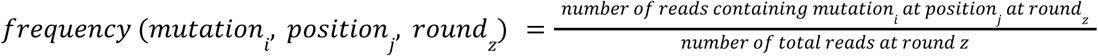

Enrichment was calculated as follows:

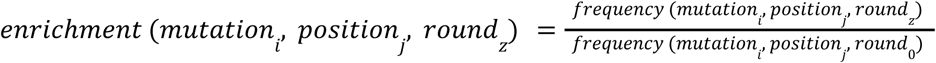

Normalized enrichment was calculated as follows:

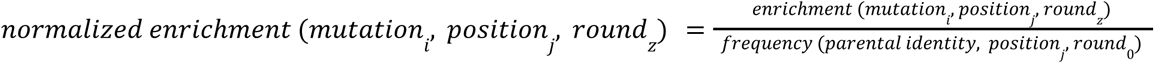

These frequencies were used to make the logoplot in Fig. 2G and the frequencies and enrichments are summarized in Supp. Table 7.

#### Mutual information analysis of D44.1 designs

For each pair of designed positions, the amino acid identity of each position for each designed sequence was added to a list one time for each read mapped to the sequence in deep sequencing of the round 3 selection. Normalized mutual information between the two positions was calculated using the normalized_mutual_info_score function in sklearn.metrics (scikit-learn v1.8.0).

#### Mutational analysis of Urelumab designs

An EpiNNet^54,55^ neural network model was trained to predict the value in each experiment for each design. In short, a multi-layer perceptron regressor with a single hidden layer the size of the number of selected positions was trained for 20,000 iterations on a one-hot encoded representation of the sequence data. Next, each pair of mutations was ranked by the difference in their predicted values for each experiment. Positions with large differences were flagged for visual inspection.

#### Prediction of MHC class II binding peptides

The light and heavy chains of each antibody were separately input to the Next-Generation IEDB Tools webserver (https://nextgen-tools.iedb.org/)^44^. The T Cell Prediction - Class II tool was used with default settings to screen the sequences against the human 27 allele panel using NetMHCIIpan 4.1 EL^61^. Hits were counted if they were in the top 10% percentile rank. The number of unique peptides for each sequence was recorded, and the number of unique peptides for the light and heavy chains for each sequence were summed.

#### Antibody numbering

Antibody numbering was performed using the AbNum^62^ web server (http://www.bioinf.org.uk/abs/abnum/). Kabat numbering^63^ is used throughout the work.

#### Distance calculations

Distance calculations were performed using PyMOL v2.3.5.

#### Data visualization

Visualization of protein structures was done using PyMOL v3.1.0. Protein surface maps were generated using the APBS^64^ plug-in in PyMOL. Data visualization was performed using matplotlib v3.1.0 and seaborn v0.9.0. Data was organized using pandas v0.24.2. Logo plots were created using Logomaker^65^.

#### Significance tests

Significance tests were performed using the ttest_ind function in SciPy v1.3.1.

### Experimental methods

#### Assembly of DNA encoding for the D44.1 library

GGAssembler^39^ was used to optimize a set of Golden-Gate^66^ overhangs to minimize the total cost of ordered oligos required to cover all mutations in the library without introducing unwanted mutations. This resulted in several variable and constant segments, with and without mutations. A gene encoding the D44.1 sequence formatted as a single-chain variable fragment was ordered from Twist BioScience. Constant segments were PCR-amplified from the gene with primers adding *BsaI* recognition sites. These and all other DNA fragments were purified using the Monarch® Spin PCR & DNA Cleanup Kit (NEB T1130). Variable segments were ordered as single-stranded degenerate oligos from IDT. The single-stranded oligos were double-stranded by a short PCR (KAPA HiFi HotStart ReadyMix; Roche 07958935001) with a single primer and purified using the Monarch kit. The concentration of each segment was measured using NanoDrop One (Thermo Scientific), and TapeStation was used to assess the purity and size of each segment. A Golden-Gate assembly was conducted using the manufacturer’s specifications for the light and heavy chains separately. Briefly, all segments are added at an equal amount, without the vector, and assembled using T4-ligase and BsaI-HF-v2 using cycles of 16°C and 37°C (New-England Biolabs, M202 and R3733, respectively). The resulting assemblies were PCR amplified using the KAPA HiFi HotStart ReadyMix (Roche Diagnostics 07958927001) and then combined using overlap-extension PCR, which also added the linker connecting the heavy and light chains. The final product was used to construct a phage-display library as described previously^67^. Briefly, the scFv fragments and library plasmid^68^ were digested with *NcoI-HF (*NEB R3193) and *NotI-HF* (NEB R3189) restriction enzymes and ligated with T4 DNA ligase (NEB M0202). Transformation of the desalted ligation into *E. coli* TG1 electro-competent cells (Lucigen 60502-1), led to 4×10^7^ transformants, greater than 1,000-fold coverage of the 7,000 variant library.

#### Phage display selections of the D44.1 library against HEWL

Panning of the phage display library against plate-coated HEWL (3 μg/ml, Sigma 62970) was carried out as described^67^. Four panning cycles were performed. The buffer was alternated between each cycle between 2% skim milk (Sigma 70166) and 2% Bovine serum albumin (Sigma A9418) in 1X PBS (Sartorius 02-023-5A) with 0.05% Tween 20 (Sigma P1379). The number of input phage at each cycle decreased (1×10^11^ for the first cycle, 5×10^10^ for the second cycle, and 1×10^10^ for the third and fourth cycles), and the number of washes increased with each cycle (from three in the first cycle to six for the fourth cycle). In an attempt to improve the k_off_ of the resultant antibodies, a parallel panning regime was carried out from the second panning cycle onward, where the last wash of the bound phages in each cycle was elongated (30, 60 and 120 minutes for the second, third and fourth cycles, respectively) and soluble HEWL (1, 1.5 and 2µM) was added.

#### Monoclonal phage ELISA

Direct ELISA was performed as described previously^69^. In short, HEWL (Sigma 62970) was used to coat (3 μg/ml) maxisorb nunc-immuno plates (Thermo 44204). Equilibrated, titrated phage were added to the blocked plate, washed, and bound phage were detected with HRP-conjugated anti-M13 antibody (Sino Biological 11973) followed by TMB substrate (Millipore ES001).

#### Antibody production

##### D44.1 and its designs

The antibodies were produced by GenScript. The amino acid sequences for the light and heavy chains were back-translated into DNA and codon optimized for expression in CHO cells using GeneSmart. The DNA was synthesized and cloned into an optimized murine IgG2a expression vector for both the light and heavy chains. The plasmids were maxiprepped to yield transfection-grade plasmids using GenScript Proprietary in-house columns and reagents. The plasmids were then transiently co-transfected at a 1:1 ratio into suspension CHO-Express cell cultures. Briefly, suspension CHO cells were grown in serum-free expression medium containing GenScript proprietary in-house buffers and supplements in Erlenmeyer Flasks at 37°C with 5% CO_2_ on an orbital shaker. One day before transfection, the cells were seeded at 3-6×10^6^ cells/mL in an Erlenmeyer flask. On the day of transfection, DNA and transfection reagent were mixed at an optimal ratio and then added to the flask with cells ready for transfection. The cell culture supernatants were collected by centrifugation at 3,000 rpm for 30 min when the cell viability was less than 50%. The supernatants were filtered and loaded onto AmMag Protein A Magnetic Beads (GenScript L00695) and incubated for the appropriate time. The beads were washed and eluted using GenScript Proprietary in-house columns and reagents. The eluted fractions were pooled and buffer exchanged to 1X PBS pH 7.2. The purified protein was analyzed by SDS-PAGE analysis using a 4%∼20% gradient SDS-PAGE gel (GenScript M42012) to determine the molecular weight and purity. Each sample was run in reducing sample buffer (60 mM Tris-HCl, 2% SDS, 6% Glycerol, 0.1% bromophenol blue, 50 mM DTT, pH 6.8) and non-reducing sample buffer (50 mM Tris-HCl, 1% SDS, 6% Glycerol, 0.1% bromophenol blue, 50 mM NEM, pH 6.8). The endotoxin level of protein was also detected using a Kinetic Turbidimetric Assay (Bioendo). The concentration was determined by measuring A280 using a NanoDrop Ultra (ThermoScientific). The final yield was calculated by multiplying the concentration by the total volume of the eluate. The final yield was then corrected by the purity measured using SDS-PAGE.

##### 6G08 and its designs for binding assays

The variable heavy and light chain sequences of the antibodies were cloned into pEE6.4 expression vectors (Lonza) containing human IgG1 constant domains. Plasmid DNA was prepared using a DNA miniprep kit (Qiagen), and constructs were verified by Sanger sequencing (Source BioScience). ExpiCHO cells (Thermo Fisher A29127) were cultured at 37°C with 8% CO2 and shaking at 125 rpm. Transient transfections were performed at a density of 6.0 X 10^6^ cells/ml following the manufacturer’s protocol, and cultures were harvested 7 days post-transfection. Antibodies were purified using a Protein A MabSelect SuRe column (Cytiva) on an AKTA Start chromatography system. Purity and integrity were confirmed by gel electrophoresis and HPLC, and endotoxin levels were measured for downstream applications.

##### 6G08 and its designs for developability assays

The antibodies were produced by GenScript. The amino acid sequences for the light and heavy chains were back-translated into DNA and codon optimized for expression in CHO cells using GeneSmart. The DNA was synthesized and cloned into an optimized human IgG1 expression vector for both the light and heavy chains. The plasmids were maxiprepped to yield transfection-grade plasmids using GenScript Proprietary in-house columns and reagents. The plasmids were then transiently co-transfected at a 1:1 ratio into suspension CHO-Express cell cultures. Briefly, suspension CHO cells were grown in serum-free expression medium containing GenScript proprietary in-house buffers and supplements in Erlenmeyer Flasks at 37°C with 5% CO_2_ on an orbital shaker. One day before transfection, the cells were seeded at 3-6×10^6^ cells/mL in an Erlenmeyer flask. On the day of transfection, DNA and transfection reagent were mixed at an optimal ratio and then added to the flask with cells ready for transfection. The cell culture supernatants were collected by centrifugation at 3,000 rpm for 30 min when the cell viability was less than 50%. The supernatants were filtered and loaded onto AmMag Protein A Magnetic Beads (GenScript L00695) and incubated for the appropriate time. The beads were washed and eluted using GenScript Proprietary in-house columns and reagents. The eluted fractions were pooled and buffer exchanged to 1X PBS pH 7.2. The endotoxin level of protein was detected using a Kinetic Turbidimetric Assay (Bioendo). The concentration was determined by measuring A280 using a NanoDrop Ultra (ThermoScientific). Analytical size-exclusion chromatography (SEC) was performed on an Waters premier UPLC system equipped with a Biozen 1.8um dsec-2 200A column (Phenomenex, 00H-4787-E0). Purified antibodies (3 µg) were loaded on the column with a mobile phase of 0.1 mol/L Na_2_SO_4_ in 0.118 mol/L Phosphate Buffer (pH 6.7±0.3) and a flow rate of 0.35 mL/min, while monitoring at 280 nm. Chromatogram traces were collected and analyzed using the Waters Empower software. Based on a single experiment.

##### Urelumab and its designs

The heavy- and light-chain variable domains of Urelumab and its designs were codon-optimized, synthesized as double-stranded DNA fragments, and assembled into human IgG1 expression vectors by Golden Gate cloning. Transient expression was performed in ExpiCHO-S cells at a viable cell density of 6 × 10^6^ cells/mL using the ExpiCHO Expression System High-Titer workflow at 3 mL scale with minor modifications (Thermo Fisher, A29133). At 18–22 h post-transfection, cultures were shifted to 32°C, 5% CO2, and supplemented with the ExpiCHO Feed/Enhancer mixture according to the manufacturer’s instructions. Supernatants were harvested on day 10 post-transfection, clarified by centrifugation and filtration through a 1.2 µm AcroPrep membrane (Cytiva, 97035). Antibodies were purified from clarified supernatants by tip-based Protein A affinity purification using MabSelect PrismA resin (IMCS, 04T-H6R80A-1-50-96). Samples were loaded by repeated aspiration–dispense cycles, washed with PBS (pH 7.2), and eluted with 0.1 M glycine-HCl (pH 2.8; Pierce, 21009), with immediate neutralization by Tris-HCl to a final pH of 7.2. Supernatant clarification and Protein A purification were executed on a Hamilton Vantage platform under fully automated control. Six low-energy and one high-energy design either did not express or had low monomer content, and were excluded from further analysis.

#### NanoDSF

##### D44.1 and its designs

The thermal stability of the antibodies was determined using the Prometheus Panta instrument (NanoTemper). Antibodies were diluted to 0.2 mg/mL in 1X PBS and the temperature was ramped from 20 °C to 100 °C, at 1.0 °C/min. The T_m_1 values were identified using the Prometheus Panta software (v. 1.4.4) with default parameters.

##### 6G08 and its designs

The NanoDSF assays were performed by GenScript. The thermal stability of the antibodies was determined using the UNcle instrument (Unchained Labs, RSP24110640). Antibodies were diluted to 1 mg/mL in 1X PBS, and the temperature was ramped from 25 °C to 95 °C, at 1.0 °C/min. The T_m_1 values were identified using the UNcle software with default parameters. Static light scattering (SLS) was also performed with four acquisitions and an acquisition time of five seconds.

##### Urelumab and its designs

The thermal stability of the antibodies was determined using the Prometheus Panta instrument (NanoTemper). Antibodies were diluted to 0.25 mg/mL in PBS pH 7.2, and the temperature was ramped from 25 °C to 95 °C at 1 °C/min. The T_m_1 values were identified using the Prometheus Panta software (v. 1.4.4) with default parameters.

#### SPR

##### D44.1 and its designs

SPR experiments on the anti-HEWL antibodies were carried out on a Biacore 8K instrument (Cytiva) at 25 °C with PBST (1X PBS + 0.05% Tween-20). For binding analysis, each antibody was immobilized to a flow cell of a protein A sensor chip (Cytiva 29127555) at a concentration of 5 μg/mL (corrected by purity from SDS-PAGE) in 1X PBS for 60 sec at a flow rate of 10 μL/min. A reference flow cell for each channel was not loaded with antibody. Approximately 100-200 response units of each antibody were immobilized. Samples of different HEWL (Sigma 62970) concentrations (10, 5, 2.5, 1.25, 0.63, 0.31, and 0.16, and 0.08 nM) were injected over the surface at a flow rate of 30 μL/min for 240 s, and the chip was washed with PBST for 600 sec at a flow rate of 30 μL/min. After each injection, surface regeneration was performed with a 30-second injection of 10 mM glycine pH 1.5 at a flow rate of 10 μL/min, and reached the same baseline. The sensogram for each flow cell without antibody was subtracted from the sensogram for the flow cell with antibody from the same channel. The acquired sensograms were analysed using the device’s software (v.4.0.8.19879). The values for k_off_ were first fit independently with a 1:1 dissociation model with k_off_ fit globally and offset and drift set to a constant value of 0. The fit values for k_off_ were then used to perform full kinetic fits to a 1:1 binding model with global fitting of *k_on_*, R_max_, and t_c_ with standard initial values. RI and drift were set to a constant 0, and k_off_ was set as a constant to the values fit previously. The experiment was repeated using different samples of HEWL and different dilutions of the antibodies. The 2.5, 1.25, 0.63, 0.31, and 0.16 nM samples were used for fitting.

##### 6G08 and its designs

SPR was performed using a Biacore T100 upgraded to T200 sensitivity (Cytiva) with a flow cell temperature set to 25°C. An anti-HIS antibody was immobilised on a Biacore CM5 chip using the His capture kit (Cytiva 28995056). Recombinant human FcγRIIb (R&D Systems) at 0.5μg/ml was captured with a target of 100 RU. Antibodies at indicated concentrations were then flowed at 30 μL/ml for 4 minutes, followed by a 5-minute dissociation with HBS-EP+ (Cytiva) used as running buffer. Data was analysed using Biacore Evaluation software and fit to a bivalent model.

##### Urelumab and its designs

SPR measurements were performed on an LSA^XT^ (Carterra) at 37°C using a capture-based non-regenerative multi-cycle kinetic experiment format. To prepare the sensor surface, the SPR system was exchanged into a running buffer of 10 mM HEPES pH 7.6, 150 mM NaCl, 3 mM EDTA, 0.05% P-20. Next, a new CMDP sensor chip (Carterra) was primed with six pulses of 200 mM Sodium Borate, 1 M NaCl, pH 9.0 (180s exposure time). Next, the surface was activated with a 1:1 mixture of 400 mM EDC and 100 mM NHS (Thermo) for 7 minutes. Next, an anti-human Fc polyclonal (Southern Biotech 2081-01) was prepared at 100 ug/mL in 10 mM Sodium Acetate pH 4.5 and injected over the surface for 7 minutes to prepare a saturated “lawn” of capture reagent. Finally, all remaining NHS-esters were deactivated by the addition of 0.1 M ethylene diamine diluted in 200 mM sodium borate pH 9.0. For measurement of binding kinetics for the antibody cohort, the system was exchanged into an experimental running buffer consisting of 10 mM HEPES pH 7.6, 150 mM NaCl, 3 mM EDTA, 0.05% P-20, 0.01% w/v BSA. Antibodies were prepared as 1 ug/mL dilution in the experimental running buffer and captured on the first quadrant using the 96-print head with an exposure time of 240s. Following this, the single flow cell was docked and a series of 10 blank injections followed by a set of 7 increasing concentrations of 4-1BB (Acro) ranging from 0.2 nM - 1000 nM (3-fold dilution) diluted into running buffer. The association phase was set to 180s followed by a dissociation phase of 600s for each blank and antigen concentration injected. Following injection of the last antigen concentration, The surface was regenerated by 3 pulses of 10 mM Glycine pH 1.7 (30s exposure time). Data was processed using a double-referencing format in Kinetics software (Carterra) and data were fit to a global 1-1 binding model. Data that significantly deviated from a 1-1 model fit or showed no discernable binding were excluded from analysis.

#### Flow cytometry staining of B-cells

Leukocyte cones were obtained from healthy donors from the Southampton NHS Blood and Transplant Service. Use was approved by the University of Southampton Faculty of Medicine Ethics Committee and East of Scotland Research Ethics Service, Research ethical committee reference number 16/ES/0048. PBMC were isolated using lymphoprep (Stemcell Technologies). 2.5×10^5^ PBMC were stained with 1 μl anti-human CD19 PE-Cy7 (Biolegend 302216) and 6G08 antibodies at the indicated concentrations for 30 minutes at 4°C. Cells were washed, and 0.5 μl PE goat anti-human IgG (Jackson Immunoresearch 109-116-098) was added for 30 minutes at 4°C. After washing, samples were run on a Cytoflex flow cytometer (Beckman Coulter). Data was analysed using FlowJo v10 (BD Biosciences) by gating on single cells, identifying lymphocytes based on FSC vs. SSC, and gating on CD19+ B cells (Supp. Fig. 10).

#### Flow cytometry staining of CHO-K1 cells overexpressing FcγRIIa

CHO-K1 (ECACC 85051005) cells stably expressing hFcγRIIa (H131) were generated as described previously^70^ and were maintained in RPMI + 10% FCS (Sigma-Aldrich), glutamine (2mM), sodium pyruvate (1mM), penicillin (100 U/ml), streptomycin (100 µg/ml) and 10 µg/ml puromycin (Invivogen), all Life Technologies unless otherwise stated. Cells were stained with 6G08 antibodies as above, washed, and run on a Cytoflex flow cytometer (Beckman Coulter). Data was analysed using FlowJo v10 (BD Biosciences) by gating on FSC vs. SSC (Supp. Fig. 10).

#### AC-SINS of 6G08 and its designs

The AC-SINS assays were performed by GenScript. AffiniPure Goat Anti-Human IgG, Fcγ fragment specific (Jackson ImmunoResearch 109-005-098) and ChromPure Goat IgG, whole molecule (Jackson ImmunoResearch 005-000-003) were each diluted to 0.4 mg/mL in 20 mM sodium acetate buffer (pH 4.3). The diluted antibodies were mixed with AuNPs (gold colloid, 20 nm, Ted Pella Inc, 15705). The mixture was gently inverted to ensure homogeneity and incubated at room temperature for at least 17 hours to allow antibody adsorption. The total coating volume was adjusted based on experimental needs, and the resulting antibody-coated AuNPs were used at 5 μL per well in a polystyrene 96-well plate (Corning 9018). A 1 mM PEG thiol stock solution (Sigma-Aldrich 729140) was diluted 100X with PBS (pH 7.2) to obtain a final concentration of 10 μM. The diluted PEG thiol solution was mixed with antibody-coated AuNPs, and the suspension was gently inverted and incubated at room temperature for one hour to block unoccupied binding sites. Following blocking, the AuNPs suspension was concentrated using a suitable syringe filter device (Cytiva 4544/4520). The filtrate was discarded, and AuNPs retained on the membrane were eluted using PBS (pH 7.2) at approximately one-tenth of the original volume or not less than 50 μL to account for membrane dead volume. Antibody-coated AuNPs and test antibody samples at a final concentration of 700 nM were added to wells of a polypropylene 96-well plate (Agilent Technologies, Inc. 5043-9311). A reference control (PBS, pH 7.2), a negative control (Daclizumab), and a positive control (Bococizumab) were included. All conditions were tested in triplicate. Plates were gently mixed and incubated at room temperature for 2 hours. Following incubation, the plate was centrifuged at 1000 rpm for 1 minute at 20 °C. Absorbance was recorded using a calibrated microplate reader (Molecular Devices iD5 RSP24110558) between 500 and 600 nm, with measurements taken every 2 nm. The wavelength shift was calculated as the difference between the maximum absorbance wavelength of the sample and the maximum absorbance wavelength of the blank control. Analysis was performed using the SoftMax Pro 7.3.1 software.

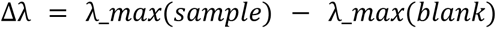

#### Ovalbumin binding of 6G08 and its designs and Urelumab and its designs

Ovalbumin binding for 6G08 and its designs was measured by Genscript using the PAIA Biotech developability kit (PA-DEV-OVA-01, PAIA Biotech). Samples were diluted to 10 µg/mL with deionized water (Bioend, TRW50S) in a 96-well round-bottom plate (Corning 9018). Fluorescently labeled nanobodies were added to a 96-deep-well plate according to the kit instructions. Sample dilution and addition were performed using the Tecan system according to the Tecan program prompts. For each well, 20 µL of sample and 40 µL of fluorescent reagent were added, with each sample in duplicates. The reaction plate, after sample addition, was sealed with a plate seal and incubated at 20 °C with orbital shaking at 2200 rpm for 30 min, followed by shaking at 1500 rpm for 10 min at 20 °C. The plate was centrifuged in a centrifuge pre-cooled to 20 °C at 1500 × g for 1 min. The plate was carefully removed after centrifugation. Fluorescence intensity was measured using a calibrated microplate reader (Molecular Devices iD5 RSP24110558) and analyzed using SoftMax Pro 7.3.1 software. Denosumab and Bevacizumab were used as positive and negative controls, respectively.

Ovalbumin binding for Urelumab and its designs was measured using the PAIA Biotech developability kit (PA-DEV-OVA; PAIA Biotech). Adalimumab and bococizumab formatted as hIgG1 were used as negative and positive controls, respectively. Samples were diluted to 10 µg/mL in ddH_2_O, then 40 µL of PAIA PSP mix and 20 µL of antibody were combined in the 384-well PA-DEV-OVA plate in duplicate. Samples were shaken for 30 mins at RT at 2200 rpm, then centrifuged at 500xg for 1 minute. Fluorescence readings were taken on an Envision plate reader using bottom read with excitation and emission at 620 and 685, respectively. Relative ovalbumin binding was calculated by normalizing the average signal between buffer only signal (no resin binding) and zero signal (full resin binding).

#### CSI-BLI of Urelumab and its designs

CSI-BLI measurements were performed on an Octet RH16 (Sartorius) with Octet Anti-Human IgG Quantitation (AHQ) Biosensors (Sartorius, Cat# 18-5001). Adalimumab and bococizumab formatted as hIgG1 were used as negative and positive controls, respectively. Test articles and controls were diluted in 1x Octet Kinetics buffer (Sartorius, Cat# 18-1105) to 3 μg/mL for loading and 150 μg/mL for association steps. Human Fc fragment (Jackson Immuno Research, 009-000-008) was diluted to 42.4 μg/mL in 1x Octet Kinetics buffer for blocking. Sensors were regenerated and neutralized using glycine pH 1.5 (Cytiva, Cat# BR100354) and 1x Octet Kinetics buffer, respectively. Samples were prepared in 384-well tilted bottom Octet plates (Sartorius, Cat# 18-5166) with 50 μL per well. Octet AHQ Biosensors were soaked in 1x Octet Kinetics buffer for at least 10 min before measurements. The experiment included baseline (60 sec in 1x Octet Kinetics buffer), loading (3 μg/mL antibodies for 100 sec), blocking (human Fc for 180 sec), baseline (60 sec in 1x Octet Kinetics buffer), association (150 μg/mL antibodies for 300 sec), dissociation (30 sec in 1x Octet Kinetics buffer), and regeneration. Data was processed in Octet Analysis Studio software with alignment to baseline and Savitzky-Golay filtering applied. CSI-BLI fold change was determined by normalizing the association response to that of the negative control, adalimumab.

#### Accelerated stability of Urelumab and its designs

Test antibodies were normalized to 1 mg/mL in PBS pH 7.2 with 10 mM EDTA and incubated for 14 days at either 4°C or 45°C in 96-well plates (Armadillo PCR Plate, Thermo Scientific AB2396). Samples were filtered with 0.2 um filters (AcroPrep Advance 96-well plates, 350 uL, wwPTFE membrane), then 5 µL was injected onto a UPLC Agilent 1290 Infinity II LC system equipped with a Waters ACQUITY Premier Protein SEC column (250Å, 1.7 µm, 150 mm). The mobile phase used was 100 mM sodium phosphate, 100 mM sodium sulfate, 0.05% w/w sodium azide, pH 6.8, with a flow rate of 0.5 mL/min. The monomer, aggregate, and fragment percentages for each antibody were determined using ChemStation software (Agilent). The change in monomer, aggregate, and fragment content was calculated from the difference between each antibody incubated at 4°C versus 45°C.

#### BSA ELISA of Urelumab and its designs

Black, 384-well high binding plates (Greiner Bio-One, Cat# 781077) were coated with blocking buffer (PBS pH 7.2 + 0.5% BSA) at 95 μL/well and incubated overnight at 4°C. After coating, plates were washed with PBS pH 7.2. Antibody samples and controls were diluted to 15 μg/mL in blocking buffer and added at 12.5 μL/well in quadruplicates. Adalimumab and bococizumab formatted as hIgG1 were used as negative and positive controls, respectively. Plates were incubated for 1 h at room temperature (RT), washed three times with PBS pH 7.2, and anti-human IgG (Fc specific)-peroxidase (Sigma-Aldrich, Cat# A0170) was added (0.3 μg/mL dilution in blocking buffer) at 12.5 μL/well. After 1 h incubation at RT, plates were washed three times with PBS pH 7.2. QuantaBlu working solution (ThermoFisher, Cat# 15169) was prepared by mixing 1 part QuantaBlu Stable Peroxide Solution with 9 parts Substrate Solution, equilibrated to RT, and added at 25 μL/well. After 20 min incubation at RT, the reaction was stopped with QuantaBlu Stop Solution at 25 μL/well. Fluorescence readings were taken on an Envision plate reader with excitation and emission at 325 nm and 420 nm, respectively. BSA scores were calculated by normalizing the average fluorescence to the average fluorescence of the positive control, bococizumab.

## Data availability

All data generated or analyzed during this study are included in this published article and its supplementary files. Amino acid sequences of the Fvs of the four D44.1, ten 6G08, and 59 Urelumab designs characterized as IgG are available in Supp. Tables 16-18. The bam files from the D44.1 deep sequencing data have been deposited in the Figshare database under accession code 32000151. The raw pod5 files can be provided upon reasonable request. The PDB files used for design are available on the Protein Data Bank with accession IDs 1MLC (D44.1), 5OCC (6G08), and 6MHR (Urelumab). Source data are provided with this paper.

## Code Availability

RosettaScripts^50^ XMLs for running LAffAb are available at https://github.com/Fleishman-Lab/LAffAb_public.

## Acknowledgments

We thank the Membrane Protein Laboratory at the Diamond Light Source for access to the Prometheus NT.48 instrument. The Membrane Protein Laboratory is funded by a grant from the Wellcome Trust (223727/Z/21/Z) with additional support provided by Diamond Light Source and the Research Complex at Harwell, both Instruct-ERIC centers. We thank Dina Listov, Olga Khersonsky, Rebecca Wilen, Inbar Cahana, Mimi Schwartz, Lucas Krauss (Weizmann Institute of Science), Ahuva Nissim (Queen Mary University of London and Weizmann Institute of Science), Shahar Harel (Technion), and Hayden Fisher (ESRF) for critical discussions on the work. A.T. was supported by Teva Pharmaceutical Industries Ltd as part of the Israeli National Forum for BioInnovators (NFBI) and by the Azrieli Foundation under the Azrieli Fellows Program. I.G.E. and M.S.C. were supported by a grant from Cancer Research UK, DRCPGM\100039. The study was supported by a European Research Council Advanced Grant (101140394) to S.J.F.

## Author Contributions

A.T. and S.J.F. conceived the idea. A.T. and S.J.F. wrote the paper, with input from all authors. A.T. developed the affinity optimization algorithm and designed all amino acid sequences. A.M. assembled the D44.1 library and performed the phage display screening. A.T. and Y.F.S. performed biophysical characterization of the anti-HEWL antibodies, and A.T., Y.F.S, S.J.F, and T.M.M. analyzed the data. S.G. prepared the anti-HEWL sequences for deep sequencing. A.T. performed the deep sequencing experiment, and A.T. and S.J.F. analyzed the data. I.T. performed the mutual information analysis. R.J.O., I.G.E., and J.K. performed experimental work related to the 6G08 designs, and A.T., S.J.F., R.O., I.G.E., and M.S.C. analyzed the data. T.P., C.D., J.B., P-H.C., and D.X.Z. performed experimental work related to the Urelumab designs, and A.T., S.J.F., T.P., T.M.M., A.B.D., and G.K. analyzed the results. O.P. and A.T. performed the mutational analysis of the Urelumab designs, and O.P., A.T., and S.J.F. analyzed the results. J.H. performed the experimental work related to the Cetuximab designs, and J.H., D.C., A.T., and S.J.F. analyzed the data. S.J.F., M.S.C., and D.C. were responsible for funding acquisition.

