## Supplementary figures and supplementary tables 1-5 and 7-12 for "Energy-guided combinatorial co-optimization of antibody affinity and stability"

Note: Supplementary Tables 6 and 13-18 are available in an accompanying Excel file.

| Position | CDR | Parental residue | Residues removed | Residues added | Final residues |
|---|---|---|---|---|---|
| $27_L$ | L1 | Q | **E** | | Q |
| $30_L$ | L1 | S | **KD** | | SGN |
| $31_L$ | L1 | N | **KT** | | N |
| $49_L$ | L2 | K | **S** | | K |
| $50_L$ | L2 | Y | **DEFHN** | | Y |
| $52_L$ | L2 | S | **T** | | S |
| $55_L$ | L2 | S | **HEF** | | SA |
| $91_L$ | L3 | S | | **T** | ST |
| $92_L$ | L3 | N | | **A** | NA |
| $93_L$ | L3 | S | **AEHY** | | S |
| $25_H$ | H1 | T | **S** | | T |
| $30_H$ | H1 | S | **KNT** | | S |
| $58_H$ | H2 | Y | **F** | | Y |

**Supplementary Table 1**. Residues added (blue) or removed (red) after visual inspection in the D44.1 library design. Amino acid numbering in all tables according to Kabat[1].

| Position | CDR | Parental residue | Residues removed | Residues added | Final residues |
|---|---|---|---|---|---|
| $26_L$ | L1 | S | **DT** | | S |
| $27_L$ | L1 | S | **DKNY** | | S |
| $48_L$ | L2 | I | **V** | | IL |
| $52_L$ | L2 | T | **K** | | T |
| $53_L$ | L2 | N | **HKRSY** | | NQ |
| $91_L$ | L3 | W | | **Y** | WY |
| $94_L$ | L3 | S | **ANY** | | S |
| $25_H$ | H1 | S | **A** | | S |
| $30_H$ | H1 | S | **A** | | S |
| $31_H$ | H1 | S | **ADHKNR** | | S |
| $33_H$ | H1 | G | **AEW** | | GPSTYV |
| $35_H$ | H1 | S | | **H** | SH |
| $50_H$ | H2 | G | **S** | | GA |
| $53_H$ | H2 | S | **ADNR** | | S |
| $54_H$ | H2 | G | **S** | | G |
| $55_H$ | H2 | G | **DEKRVHNS** | | GTA |
| $95_H$ | H3 | S | **V** | | S |
| $97_H$ | H3 | G | **Y** | | GAS |
| $98_H$ | H3 | A | **DS** | | ALRYG |
| $100D_H$ | H3 | F | | **L** | FL |
| $102_H$ | H3 | I | | **V** | IV |

**Supplementary Table 2**. Residues added (blue) or removed (red) after visual inspection in the 6G08 design process.

| Position | CDR | Parental residue | Residues removed | Residues added | Final residues |
|---|---|---|---|---|---|
| $27_L$ | L1 | Q | **EH** | | QKR |
| $30_L$ | L1 | S | **KDN** | | SG |
| $31_L$ | L1 | S | **HINY** | | SKRT |
| $32_L$ | L1 | Y | **F** | | Y |
| $53_L$ | L2 | N | **IKRY** | | NA |
| $90_L$ | L3 | Q | **E** | | Q |
| $26_H$ | H1 | G | **ADS** | | G |
| $30_H$ | H1 | S | **NT** | | SKR |
| $31_H$ | H1 | G | **DEHN** | | GAS |
| $32_H$ | H1 | Y | **F** | | Y |
| $53_H$ | H2 | H | **DN** | | HFY |
| $54_H$ | H2 | G | **DKNR** | | GAS |
| $100_H$ | H3 | N | **DLR** | | NAS |
| $100C_H$ | H3 | D | **LRS** | | DANTY |

**Supplementary Table 3**. Residues added (blue) or removed (red) after visual inspection in the Urelumab design process.

| CDR | Definition used in this work | CUMAb[2] definition | Kabat[1] definition |
|---|---|---|---|
| L1 | 24-34 | 24-34 | 24-34 |
| L2 | 46-55 | 46-55 | 50-56 |
| L3 | 89-97 | 89-97 | 89-97 |
| H1 | 24-35B | 24-35B | 31-35B |
| H2 | 50-58 | 47-58 | 50-65 |
| H3 | 95-102 | 93-102 | 95-102 |

**Supplementary Table 4**. CDR definitions used in this work compared to CUMAb[2] and Kabat[1].

| Colony number | $30_L$ | $51_L$ | $54_L$ | $55_L$ | $91_L$ | $54_H$ | $56_H$ | $98_H$ | $101_H$ |
|---|---|---|---|---|---|---|---|---|---|
| **D44.1** | **S** | **V** | **S** | **S** | **S** | **S** | **S** | **N** | **G** |
| 1 | G | G | S | A | S | S | S | Y | G |
| 2 | G | G | L | A | S | S | S | Y | G |
| 3 | G | G | L | A | S | T | S | S | H |
| 4 | G | G | S | S | S | S | V | Y | G |
| 5 | G | G | S | A | S | T | V | Y | G |
| 6 (Des2) | G | G | S | A | S | S | V | Y | H |
| 7 | G | T | L | A | S | T | V | A | H |
| 8 (Des4) | G | G | S | S | S | T | V | S | H |
| 9 | G | G | S | A | S | S | V | A | H |
| 10 (Des3) | G | T | L | A | S | T | V | Y | G |
| 11 | G | T | S | A | S | S | S | S | H |
| 12 | N | T | L | A | S | S | V | Y | G |
| 13 | G | G | S | A | S | T | S | Y | G |
| 14 | G | G | L | S | S | T | V | Y | G |
| deletion in light chain | G | G | L | A | S | S | N | A | G |
| 15 | G | G | L | A | S | S | S | S | H |
| 16 | S | G | S | A | S | S | V | Y | G |
| 17 | G | G | S | A | S | S | S | A | G |
| 18 (Des1) | G | G | L | A | S | S | S | A | H |
| 19 | S | G | L | A | T | S | V | S | H |

**Supplementary Table 5**. Amino acid identities at designed positions in selected colonies of LAffAb designs based on D44.1.

| Mutation | Fraction of reads before selection | Fraction of reads after 3 selection rounds | Enrichment | Normalized enrichment |
|---|---|---|---|---|
| $S30_LS$ | 0.21 | 0.09 | 0.45 | 1 |
| $S30_LG$ | 0.68 | 0.85 | 1.25 | 2.8 |
| $S30_LN$ | 0.11 | 0.05 | 0.48 | 1.07 |
| $V51_LV$ | 0.19 | 0.03 | 0.17 | 1 |
| $V51_LT$ | 0.58 | 0.32 | 0.55 | 3.24 |
| $V51_LG$ | 0.22 | 0.64 | 2.87 | 16.79 |
| $S54_LS$ | 0.47 | 0.27 | 0.58 | 1 |
| $S54_LL$ | 0.53 | 0.73 | 1.37 | 2.34 |
| $S55_LS$ | 0.44 | 0.27 | 0.61 | 1 |
| $S55_LA$ | 0.56 | 0.73 | 1.3 | 2.12 |
| $S91_LS$ | 0.54 | 0.97 | 1.8 | 1 |
| $S91_LT$ | 0.46 | 0.03 | 0.07 | 0.04 |
| $S54_HS$ | 0.39 | 0.63 | 1.63 | 1 |
| $S54_HN$ | 0.31 | 0.03 | 0.1 | 0.06 |
| $S54_HT$ | 0.3 | 0.34 | 1.12 | 0.69 |
| $S56_HS$ | 0.21 | 0.33 | 1.52 | 1 |
| $S56_HV$ | 0.32 | 0.43 | 1.36 | 0.89 |
| $S56_HN$ | 0.22 | 0.18 | 0.82 | 0.54 |
| $S56_HI$ | 0.25 | 0.07 | 0.26 | 0.17 |
| $N98_HN$ | 0.15 | 0.03 | 0.18 | 1 |
| $N98_HA$ | 0.36 | 0.32 | 0.88 | 4.8 |
| $N98_HS$ | 0.31 | 0.29 | 0.92 | 5.01 |
| $N98_HY$ | 0.17 | 0.37 | 2.12 | 11.61 |
| $G101_HG$ | 0.65 | 0.45 | 0.7 | 1 |
| $G101_HH$ | 0.35 | 0.55 | 1.55 | 2.2 |

**Supplementary Table 7**. Statistics from deep sequencing of designed mutations in the D44.1 library.

| Mutation | Cα distance (Å) |
|---|---|
| $S30_LG$ | 0.02 |
| $S30_LN$ | 0.06 |
| $V51_LT$ | 0.05 |
| $V51_LG$ | 0.11 |
| $S54_LL$ | 0.29 |
| $S55_LA$ | 0.06 |
| $S91_LT$ | 0.13 |
| $S54_HN$ | 0.41 |
| $S54_HT$ | 0.07 |
| $S56_HV$ | 0.35 |
| $S56_HN$ | 0.46 |
| $S56_HI$ | 0.65 |
| $N98_HA$ | 0.07 |
| $N98_HS$ | 0.03 |
| $N98_HY$ | 0.05 |
| $G101_HH$ | 0.04 |

**Supplementary Table 8**. Cα distances between the design model and relaxed parental structure for each mutation in the designed D44.1 library.

| Design | $48_L$ | $53_L$ | $91_L$ | $28_H$ | $33_H$ | $35_H$ | $52A_H$ | $55_H$ | $97_H$ | $98_H$ | $100_H$ | $100D_H$ | $102_H$ |
|---|---|---|---|---|---|---|---|---|---|---|---|---|---|
| **6G08** | **I** | **N** | **W** | **T** | **G** | **S** | **G** | **G** | **G** | **A** | **A** | **F** | **I** |
| Des1 | L | N | W | N | Y | H | G | A | G | G | A | F | I |
| Des2 | L | N | W | T | T | H | G | T | S | A | A | F | V |
| Des3 | L | Q | Y | N | Y | H | G | G | G | A | A | F | I |
| Des4 | L | N | Y | T | Y | S | A | T | G | G | A | F | I |
| Des5 | L | N | W | N | Y | H | A | G | G | L | A | F | I |
| Des6 | L | Q | W | T | P | S | A | T | G | R | A | F | I |
| Des7 | I | Q | W | N | S | H | G | G | G | Y | A | L | I |
| Des8 | L | N | Y | T | S | H | G | T | G | L | A | F | I |
| Des9 | L | N | W | T | T | H | G | G | S | L | S | F | I |
| Des10 | L | N | W | N | V | H | A | A | G | A | A | F | I |

**Supplementary Table 9**. Mutations at each designed position for the ten 6G08 designs.

| Design | Number of peptides MHCII |
|---|---|
| 6G08 | 23 |
| Des1 | 24 |
| Des2 | 24 |
| Des3 | 23 |
| Des4 | 24 |
| Des5 | 23 |
| Des6 | 24 |
| Des7 | 23 |
| Des8 | 23 |
| Des9 | 22 |
| Des10 | 24 |
| Pembrolizumab | 23 |

**Supplementary Table 10**. Number of peptides predicted to bind a panel of 27 human MHC class II alleles for the ten 6G08 designs.

| Mutation | Cα distance |
|---|---|
| $I48_LL$ | 0.03 |
| $N53_LQ$ | 0.01 |
| $W91_LY$ | 0.07 |
| $T28_HN$ | 0.05 |
| $G33_HY$ | 0.33 |
| $G33_HT$ | 0.08 |
| $G33_HP$ | 0.02 |
| $G33_HS$ | 0.04 |
| $G33_HV$ | 0.26 |
| $S35_HH$ | 0.15 |
| $G52A_HA$ | 0.42 |
| $G55_HA$ | 0.04 |
| $G55_HT$ | 0.08 |
| $G97_HS$ | 0.21 |
| $A98_HG$ | 0.03 |
| $A98_HL$ | 0.04 |
| $A98_HR$ | 0.03 |
| $A98_HY$ | 0.10 |
| $A100_HS$ | 0.03 |
| $F100D_HL$ | 0.12 |
| $I102_HV$ | 0.02 |

**Supplementary Table 11**. Cα distances between the design model and relaxed parental structure for each mutation in the ten 6G08 designs.

| Design | Yield (mg/mL culture volume) | Tm1 (°C) | Tagg (°C) | SEC % monomer | AC-SINS Δλmax (nm) | PAIA Ova-Fc FI (AU) |
|---|---|---|---|---|---|---|
| 6G08 | 0.593 | 69.24 | 73.50 | 92 | 0.67 | 53360.5 |
| Des4 | 0.76 | 69.71 | 74.30 | 94 | 0.67 | 53687 |
| Des8 | 0.69 | 69.38 | 74.01 | 96 | 1.33 | 53249.5 |
| Bococizumab | n/a | n/a | n/a | n/a | 16.67 | n/a |
| Daclizumab | n/a | n/a | n/a | n/a | 1.33 | n/a |
| Denosumab | n/a | n/a | n/a | n/a | n/a | 23760.5 |
| Bevacizumab | n/a | n/a | n/a | n/a | n/a | 46233 |

**Supplementary Table 12**. Developability assays for 6G08, its designs, and therapeutic controls.

### Supplementary Figures:

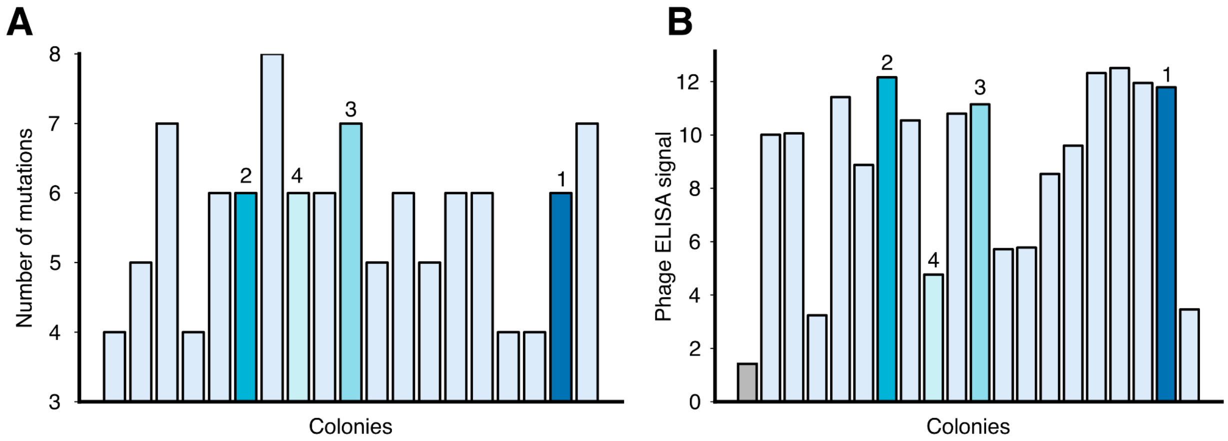


**Supplementary Figure 1. Colonies selected after four rounds of panning the D44.1 library against HEWL.** (**A**) Number of mutations away from the parental antibody for each of the 19 unique colonies sequenced after the fourth selection round. (**B**) The 19 colonies were assayed for HEWL binding using phage ELISA. Parental antibody in gray. Shown is a representative experiment out of two repetitions. (**A, B**) The four designs assayed as IgG2a are colored; the design numbers are denoted above the bars.

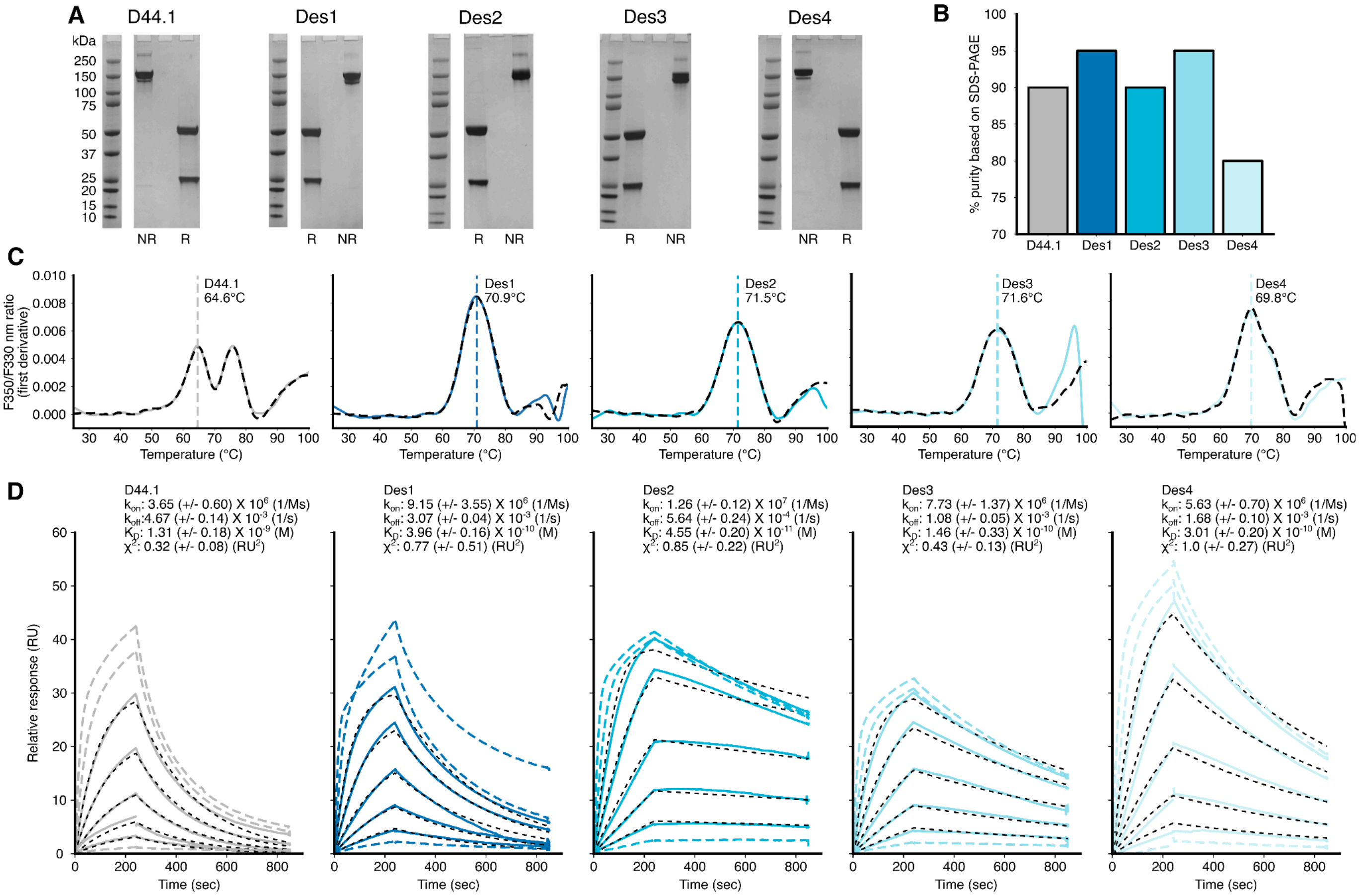


**Supplementary Figure 2. Biophysical characterization of the D44.1 designs formatted as IgG2A.** (**A**) SDS-PAGE gels of D44.1 and the four designs. Marker shown on the left. R refers to samples run in reducing sample buffer, and NR refers to samples run in non-reducing sample buffer. (**B**) Quantification of the purity of D44.1 and the four designs based on SDS-PAGE gels in A. Based on a single measurement. (**C**) Thermal denaturation of D44.1 and the four designs using nanoDSF. In each graph, the vertical, colored dashed line represents the mean of the first apparent melting temperature of two technical replicates. The colored plot represents one measurement, and the black, dashed plot is of a technical replicate. (**D**) SPR kinetic analysis of D44.1 and the four designs. Antibodies were immobilized on a protein A chip. HEWL was then assayed at 10, 5, 2.5, 1.25, 0.63, 0.31, 0.16, and 0.08 nM. Representative sensogram is shown for each antibody. Solid lines are the HEWL concentration used for fitting, and black dashed lines are the fits. Colored dashed lines are concentrations not used in fits. Values reported are the means +/- standard deviations of two technical replicates.

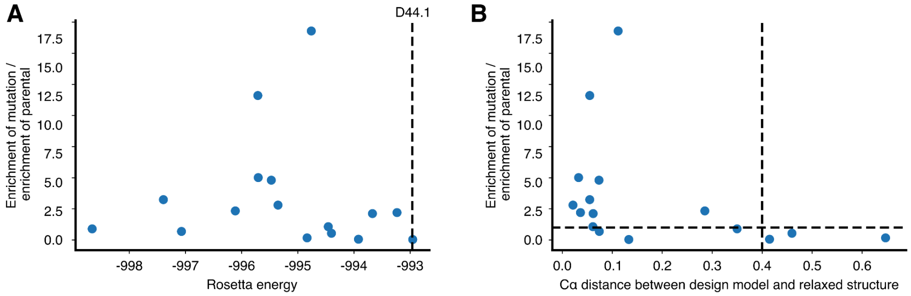


**Supplementary Figure 3: Calculated parameters vs. enrichment of individual mutations in the D44.1 library.** (**A**) Fold-change of enrichment of each designed mutation compared to the enrichment of the parental identity vs. Rosetta energy. The black dashed line represents the Rosetta energy of D44.1. (**B**) Fold-change of enrichment of each designed mutation compared to the enrichment of the parental identity vs. the Cα distance between the design model and relaxed structure. The horizontal black dashed line represents an enrichment value of 1. The vertical black dashed line represents a potential cutoff that could be used in future design projects.

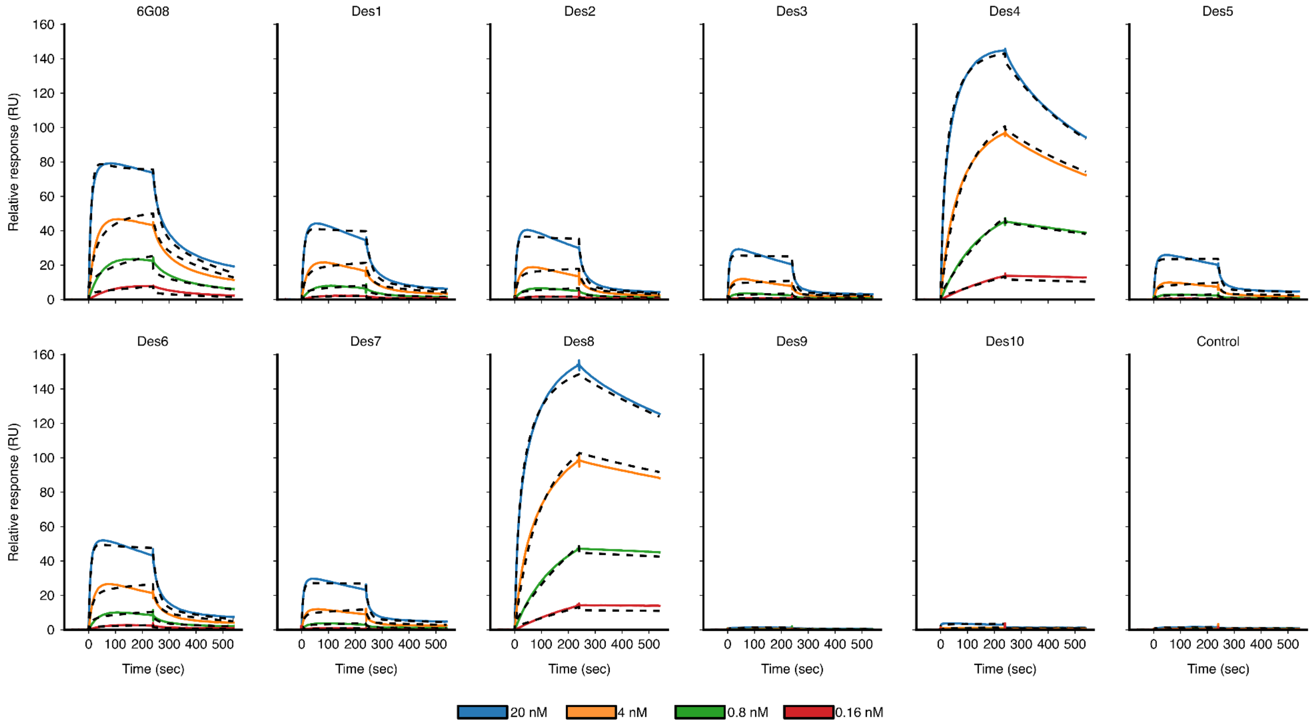


**Supplementary Figure 4: Affinity characterization of 6G08 designs formatted as human IgG1.** SPR kinetic analysis of 6G08 and the ten designs. FcγRIIb was immobilized on an anti-His chip. Antibodies were assayed at 20, 4, 0.8, and 0.16 nM. Based on a single measurement.

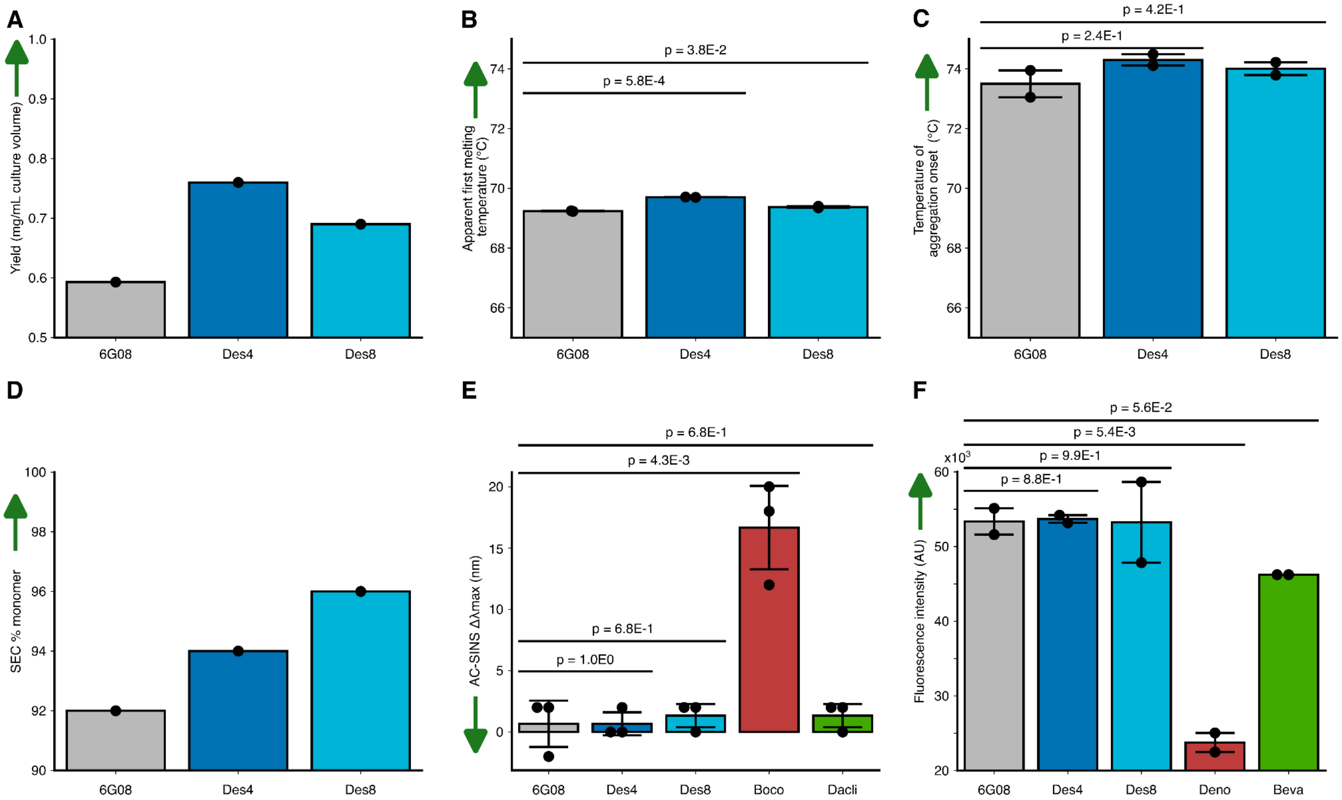


**Supplementary Figure 5: Developability characterization of 6G08 designs formatted as human IgG1.** (**A**) Expression yield**.** Based on a single measurement. (**B**) Thermal stability measured by NanoDSF. Based on two technical replicates. (**C**) Thermal aggregation resistance measured by Static Light Scattering (SLS). Based on two technical replicates. (**D**) Percent monomer measured by SEC. Based on a single measurement. (**E**) Self-association measured by Affinity-Capture Self-Interaction Nanoparticle Spectroscopy (AC-SINS). Bococizumab (Boco) and Daclizumab (Dacli) are therapeutic antibodies known to perform poorly and well in this assay, respectively. Based on three technical replicates. (**F**) Polyspecificity measured using the PAIA OVA-Fc kit. Denosumab (Deno) and Bevacizumab (Beva) are therapeutic antibodies known to behave poorly and well in this assay, respectively. Based on two technical replicates. (**A-F**) Bars and error bars represent the mean and standard deviation of technical replicates (measurements plotted as dots). Green arrows represent the desirable direction in each assay. Experiments performed by Genscript. (**B, C, E, F**) *P*-values calculated using a Student's *t*-test.

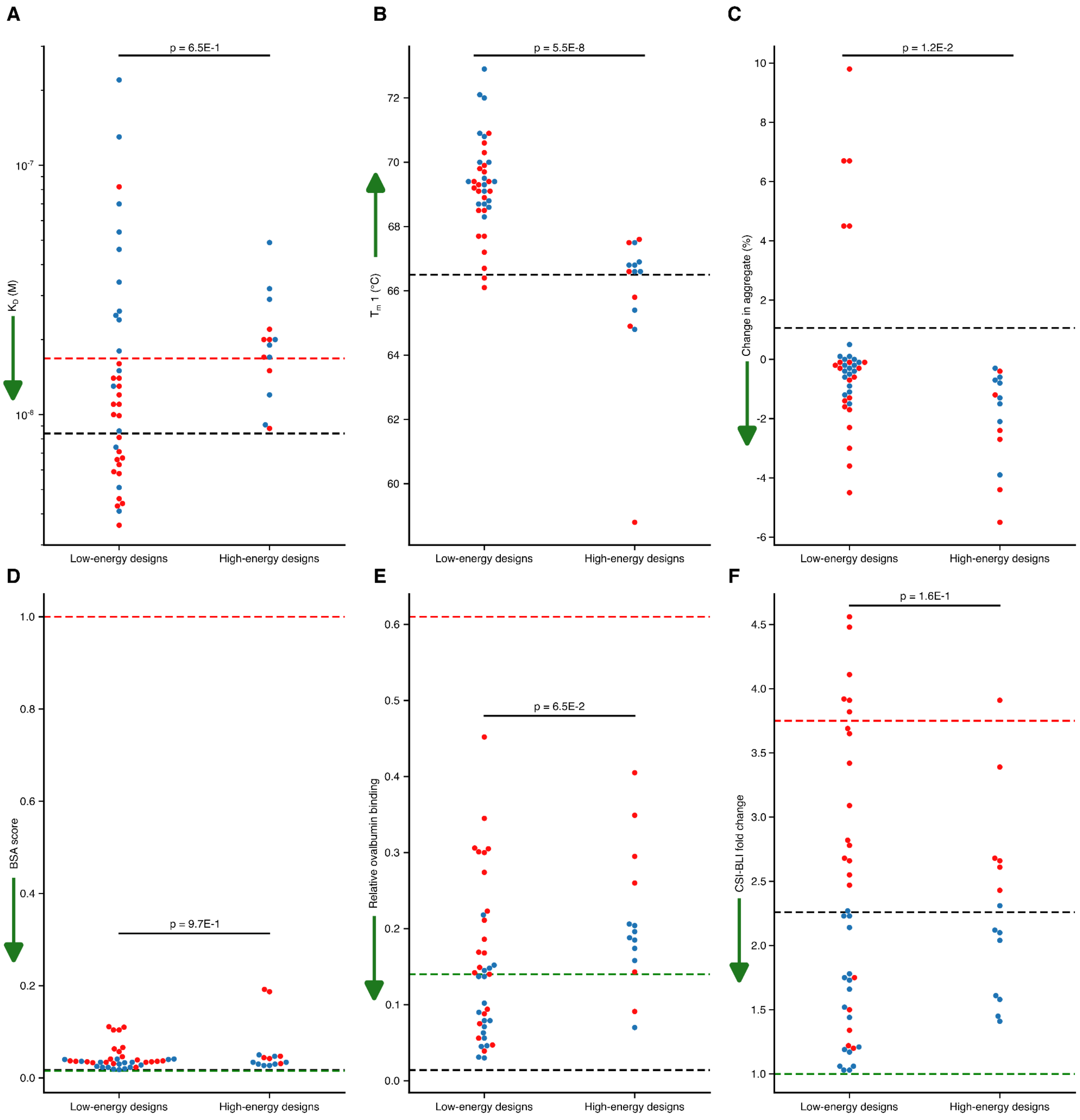


**Supplementary Figure 6: Biophysical assays of low- and high-energy designs of Urelumab.** (**A**) Affinity measured by SPR. (**B**) Thermal stability measured by nanoDSF. (**C**) Aggregation resistance in an accelerated stability assay. (**D**) Nonspecific binding measured by binding to BSA. (**E**) Nonspecific binding measured by binding to ovalbumin. (**F**) Reversible self-association measured using antibody clone self-interaction (CSI) using bio-layer interferometry (BLI)[3]. (**A-F**) Blue and red dots represent designs categorized as developable and non-developable, respectively. The black dashed line represents the value for Urelumab. *P*-values calculated using a Student's *t*-test. Dots represent the means of three technical repeats. Green arrows represent the desirable direction in each assay. (**D-F**) Green and red dashed lines represent the values for Adalimumab and Bococizumab, respectively.

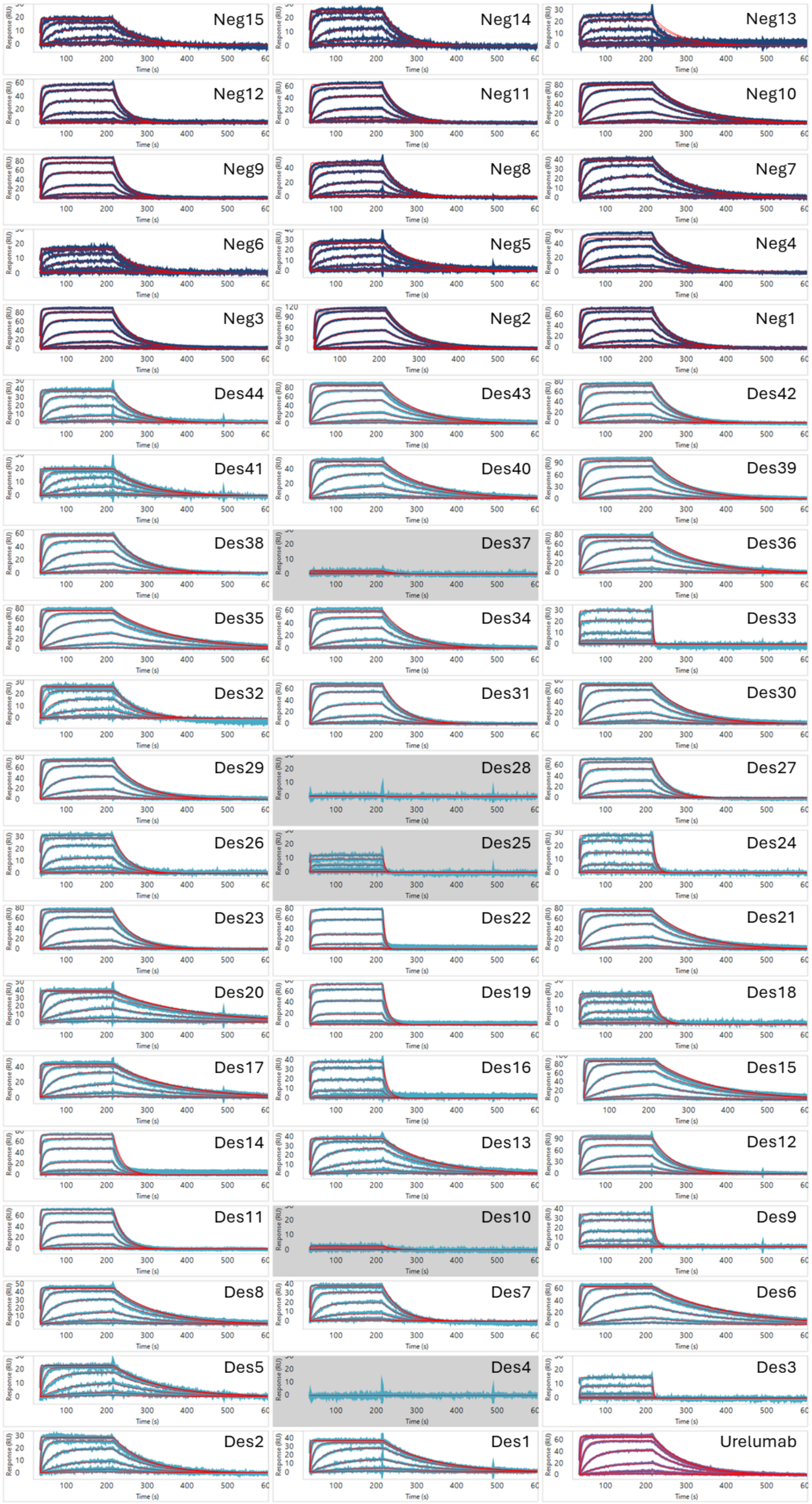

**Supplementary Figure 7: Representative sensorgrams for kinetics measurements of Urelumab and its designs.** Designs labeled Neg are high-energy designs.

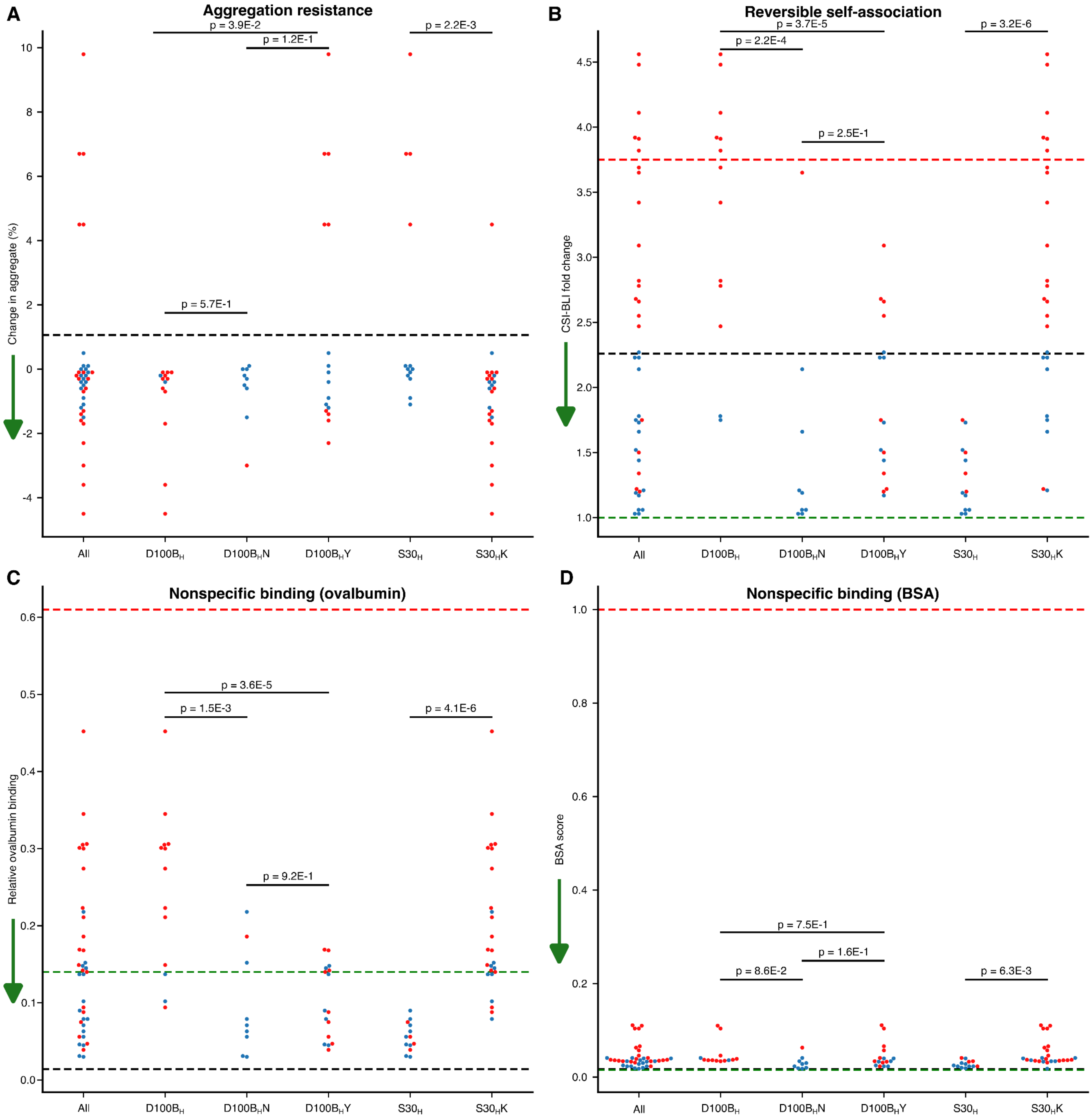


**Supplementary Figure 8: Developability assays for Urelumab and its designs.** (**A**) Aggregation resistance in an accelerated stability assay. (**B**) Nonspecific binding measured by binding to BSA. (**C**) Nonspecific binding measured by binding to ovalbumin. (**D**) Reversible self-association measured using CSI-BLI. (**A-D**) Blue and red dots represent designs categorized as developable and non-developable, respectively. The black dashed line represents the value for Urelumab. *P*-values calculated using a Student's *t*-test. Dots represent the means of three technical repeats. Green arrows represent the desirable direction in each assay. (**B-D**) Green and red dashed lines represent the values for Adalimumab and Bococizumab, respectively.

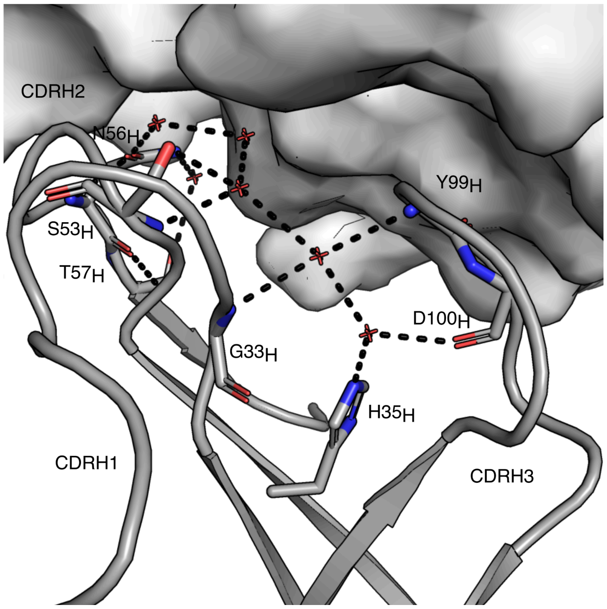


**Supplementary Figure 9: A network of water molecules supports the conformation of CDR H3 in Cetuximab.** Crystal structure of Cetuximab (PDB ID 1YY9[4]. EGFR is shown in a white surface map, and Cetuximab is shown in a gray cartoon. Key interactions with the water molecules shown in sticks.

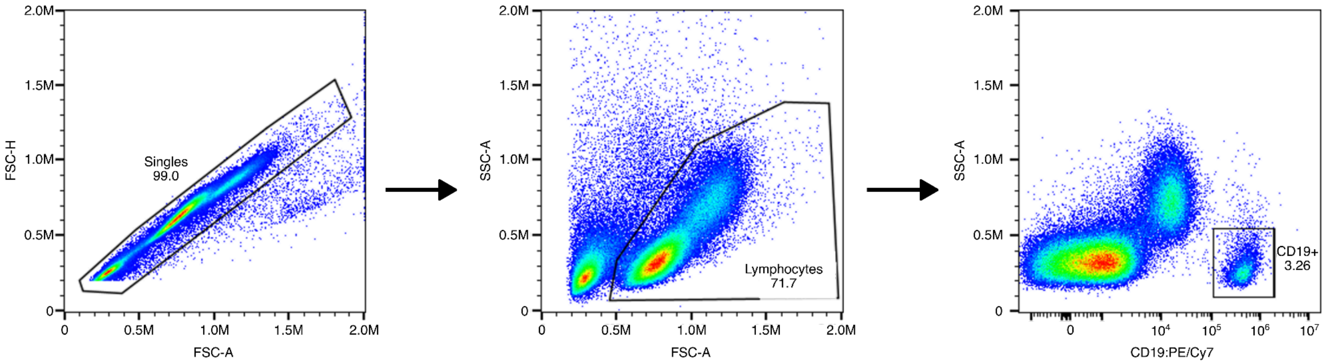


**Supplementary Figure 10: FACS gating strategy for 6G08 experiments.**
